# Proteostasis Stress Drives Stem Cell Aging, Clonal Hematopoiesis and Leukemia

**DOI:** 10.1101/2025.11.27.690982

**Authors:** Fanny J. Zhou, Michelle K. Le, Helen C. Wang, Wei Yang, Huan-You Wang, Arukshita Tiwari, Xinjian Cen, Mary Jean Sunshine, Jeffrey A. Magee, Robert A.J. Signer

**Affiliations:** Division of Regenerative Medicine, Department of Medicine, Stem Cell Discovery Center, Sanford Stem Cell Institute, Moores Cancer Center, University of California San Diego; La Jolla, CA, 92037, USA; Department of Pediatrics, Division of Hematology and Oncology, Washington University School of Medicine; St. Louis, MO 63110, USA; Department of Genetics, Washington University School of Medicine; St. Louis, MO 63110, USA; Division of Laboratory and Genomic Medicine, Department of Pathology, University of California San Diego Health Sciences; La Jolla, CA 92037, USA; La Jolla Institute for Immunology; La Jolla, CA 92037, USA

## Abstract

Aging is the primary risk factor for clonal hematopoiesis and the development of hematologic malignancies (*1–5*), yet the selective pressures that shape stem cell behavior and clonal expansion during aging remain poorly defined. Here, we identify proteostasis stress as a central driver of hematopoietic stem cell (HSC) aging and clonal evolution. We show that Heat shock factor 1 (Hsf1) is activated in aging HSCs to preserve proteostasis and sustain self-renewal. However, this physiological, age-associated adaptive mechanism is co-opted by pre-leukemic *Dnmt3a*-mutant HSCs to resist proteostasis and inflammatory stress required to fuel clonal expansion during aging. In the context of co-occurring *Dnmt3a* and *Nras* mutations, which are frequently observed in human acute myeloid leukemia (AML) (*6–13*), mutant HSCs and progenitors exhibit heightened dependence on Hsf1 for expansion, malignant transformation and disease progression. Loss of *Hsf1*, or disruption of proteostasis, impairs expansion of mutant progenitors, delays leukemia onset, and prolongs survival. Together, these findings reveal proteostasis as a key constraint in the aging hematopoietic system that imposes a selective bottleneck. Hsf1 activation enables both physiological adaptation in aging stem cells and pathological clonal outgrowth in pre-leukemic and leukemic states, establishing proteostasis control as a pivotal mechanism linking stem cell aging to clonal hematopoiesis and malignancy.

## Main Text

Aging is the single greatest risk factor for the development of cancer (*14–18*), yet the biological mechanisms that drive this increased susceptibility remain poorly understood. Over half of all blood cancers, including leukemias, myelodysplastic syndromes and myeloproliferative neoplasms, arise in individuals over the age of 60, and these malignancies are often more aggressive and harder to treat with poorer prognosis due to both high-risk genetic profiles and the decreased tolerance of older patients to intensive therapies (*19–21*). Thus, uncovering age-associated mechanisms that drive clonal hematopoiesis and malignant transformation, and that may be selectively targeted with minimal toxicity, remains an urgent clinical priority.

Although the gradual accumulation of somatic mutations has long been viewed as the primary driver of age-related cancer (*2, 22, 23*), mounting evidence suggests that broader age-associated cellular and tissue-level changes alter selective pressures in ways that promote transformation. Hallmarks of aging, including mitochondrial dysfunction, cellular senescence, epigenetic alterations and chronic inflammation can apply selective pressures, foster permissive environments, or impose bottlenecks that select for clones with oncogenic mutations that may ultimately lead to cancer (*16, 17, 24–27*). This paradigm suggests that cancer is not simply a genetic or epigenetic disorder; it instead reflects a culmination of widespread cellular changes that accrue with age.

Consistent with this model, many high-risk leukemogenic mutations arise disproportionately in older adults. Mutations in genes such as *DNMT3A* (*7, 28–30*), *TET2* (*31, 32*), *SRSF2* (*33*), *IDH1/2* (*10, 34, 35*) and *RUNX1* (*36, 37*) occur more commonly in older adults with acute myeloid leukemia (AML) than in younger individuals (*38, 39*), suggesting that these mutations confer a stronger selective advantage to pre-leukemic hematopoietic stem cells (HSCs) or other progenitors in the context of aging than they do at earlier stages of life. In support of this idea, many of these same mutations frequently occur in age-related clonal hematopoiesis (*40, 41*), a pre-malignant state characterized by the expansion of mutant HSCs that is associated with increased risk of hematologic malignancy, cardiovascular disease, and all-cause mortality (*28, 30, 42–49*). Despite extensive genetic characterization, the nature of the selective pressures imposed by aging that favor the emergence of these mutant clones remains incompletely defined.

Aging profoundly alters HSC biology (*50*). Aged HSCs exhibit functional changes, including impaired regenerative capacity, diminished self-renewal, and myeloid-biased differentiation (*51–58*). These phenotypes are accompanied by canonical hallmarks of aging such as genomic instability, telomere attrition, epigenetic reprogramming, and mitochondrial dysfunction, each of which can apply selective pressures on HSCs (*59–72*). DNA damage or inflammation can promote HSC activation and differentiation (*64, 73–79*). Changes in DNA methylation, histone acetylation/methylation or mitochondrial activity can promote HSC exhaustion (*54, 62, 80–87*). Tumor suppressor genes can be activated in response to these stressors to promote senescence and apoptosis (*70*). These processes can all reduce clonal diversity within the HSC pool and select for mutations that bypass or buffer these stressors, providing a pronounced fitness advantage to aging HSCs.

Loss of protein homeostasis (proteostasis) is one of the least understood hallmarks of aging (*16, 17*), particularly as it relates to both stem cell aging and malignant transformation. As an organism ages, misfolded proteins can accumulate in post-mitotic cells, or in cells that are largely quiescent (*88–95*). HSCs are largely quiescent (*63, 96–98*) and particularly susceptible to a loss of proteostasis (*99*). Adult HSCs maintain unusually low rates of protein synthesis relative to lineage-committed blood progenitors (*100*). Low protein synthesis helps HSCs maintain proteostasis by reducing the biogenesis of misfolded proteins and is essential for preserving long-term self-renewal capacity (*100–102*). Disruption of proteostasis can trigger HSC division, stress responses, and functional decline (*101, 103–106*). These observations raise the possibility that proteostasis pathways may serve as previously unrecognized selective pressures in the aging HSC compartment.

Here, we show that the proteostasis regulator Heat shock factor 1 (Hsf1) (*107–109*) is activated in aging HSCs and orchestrates a transcriptional program that preserves proteome integrity and stem cell function. While this adaptive response maintains HSC fitness during physiological aging, it is co-opted by pre-leukemic clones to resist cellular stress and fuel clonal expansion. Our findings identify proteostasis as a critical node at the interface between aging, clonal hematopoiesis and leukemogenesis.

### Proteostasis is transcriptionally rewired in aged HSCs

To uncover intrinsic transcriptional changes in HSCs during aging, we performed RNA sequencing on CD150⁺CD48⁻Lineage⁻Sca1⁺c-Kit⁺ (CD150⁺CD48⁻LSK) HSCs (*110*) isolated from young (3-4-month-old) and old (22-24-month-old) wild-type (WT) C57BL/6 mice (fig. S1A). Differential gene expression analysis revealed that 706 genes were altered ≥ 2-fold (p_adj_ < 0.05; fig. S1B), consistent with established transcriptional aging signatures in mouse HSCs (*111*) (fig. S1C).

Gene set enrichment analysis (GSEA) (*112*) revealed striking upregulation of pathways related to proteostasis in aged HSCs, including “Protein Targeting”, “Cytoplasmic Translation”, “Protein Folding” and “Ribosome Biogenesis” (Fig. 1A, table S1). Aged HSCs downregulate pathways related to “DNA Replication,” “Cell Division,” and “DNA Repair,” consistent with previous studies (*54, 62*) (fig. S1D, table S2). These findings suggested a robust transcriptional activation of proteostasis machinery during HSC aging. However, in surprising contrast to these widespread transcriptional changes, we observed no significant change in global protein synthesis rates (*100*) or unfolded (*113*) and misfolded protein (*114*) abundance in old versus young HSCs (Fig. 1B-D), suggesting that proteostasis is largely maintained in aging HSCs.

**Figure 1.**
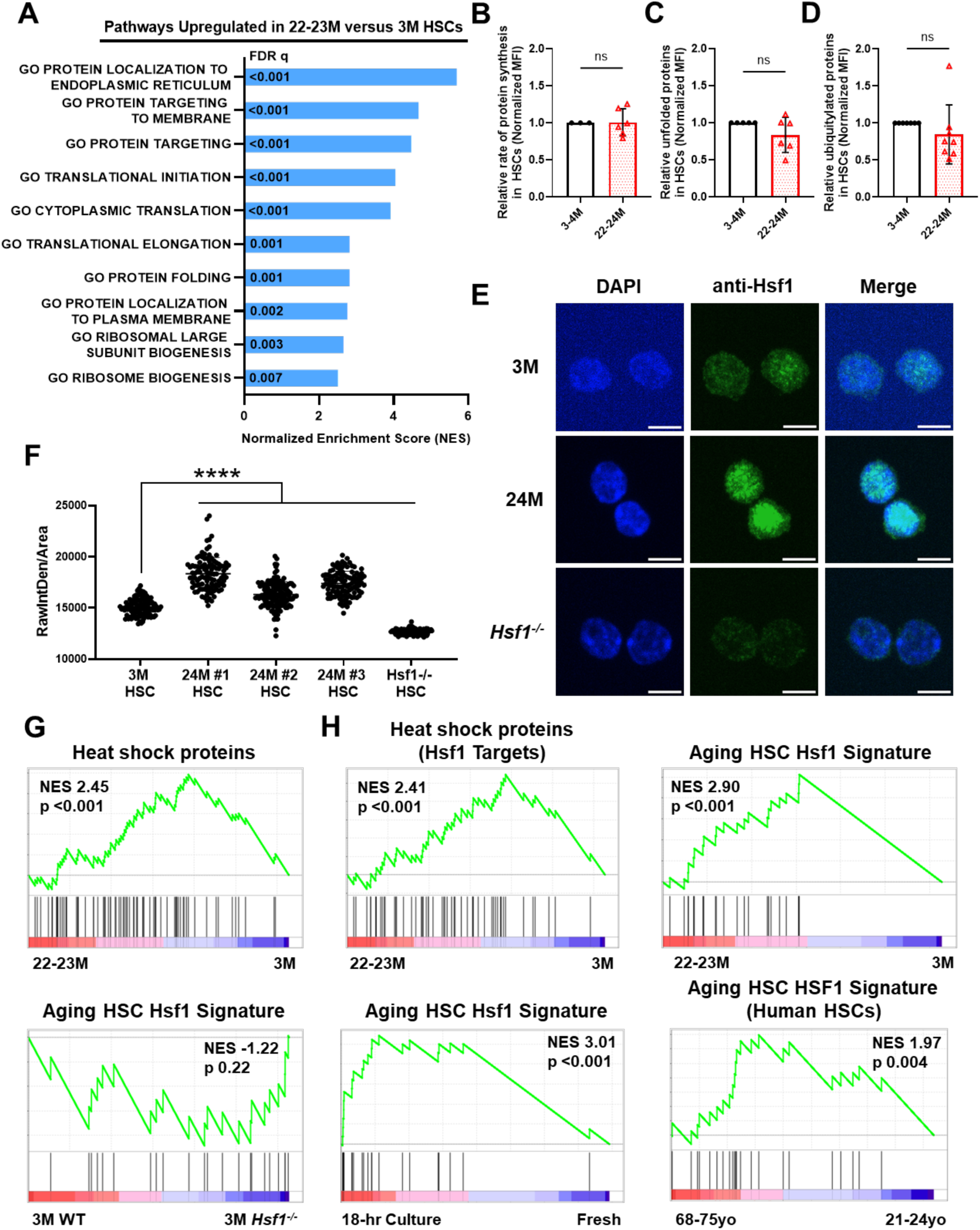
Proteostasis is transcriptionally rewired in aged HSCs. **(A)** GSEA in 3- and 22-23-month-old CD150^+^CD48^−^LSK HSCs from RNA-sequencing. Selected upregulated gene sets are shown with overlay of false discovery rates (FDR q; n=3 mice/age). **(B-D)** Relative (B) rates of protein synthesis (based on o-propargyl-puromycin incorporation), **(C)** unfolded protein abundance (based on tetraphenylethene maleimide fluorescence) and (D) misfolded protein abundance (based on polyubiquitylated protein) in 3-4- and 22-24-month-old HSCs (n=3-8 mice/age). **(E,F)** Representative immunofluorescence images (E) and nuclear quantification (F) from 3- and 24-month-old WT and 24-month-old *Hsf1^-/-^* HSCs stained with DAPI and anti-Hsf1 (Alexa Fluor 488). Scale bars represent 5μm. Each dot in (F) represents an individual HSC. **(G)** Gene set enrichment plot showing upregulation of heat shock protein (HSP) genes in 22-23-versus 3-month-old HSCs (n=3 mice/age). **(H)** Gene set enrichment plot showing upregulation of previously verified Hsf1-target HSP genes (based on HSF1 Base) in 22-23- versus 3-month-old HSCs (n=3 mice/age). **(I)** Gene set enrichment plot showing the “Aging HSC Hsf1” signature in 22-23-versus 3-month-old HSCs (n=3 mice/age). **(J)** Gene set enrichment plot showing that the “Aging HSC Hsf1” signature is not enriched in 3-month-old WT versus *Hsf1^-/-^*HSCs (n=3 mice/genotype). **(K)** Gene set enrichment plot showing upregulation of the “Aging HSC Hsf1” signature in 18-hour cultured young as compared to freshly isolated HSCs (from GSE179412, n=3 mice/condition). **(L)** Gene set enrichment plot showing upregulation of the “Aging HSC Hsf1” signature in 68-75 year-old human HSCs (from GSE188889, n=3 donors/age). Data represent mean ± SD in (B-D,F). Statistical significance was assessed using unpaired Student’s *t-*test in (B-D) and ordinary one-way ANOVA followed by Dunnett’s test relative to 3-month-old HSCs in (F).

This apparent disconnect is paradoxical: why are proteostasis programs transcriptionally upregulated if proteostasis appears largely intact? Prior work has shown that HSC function is exquisitely sensitive to proteostasis disruption, with even modest imbalances impairing self-renewal (*100, 101*). Therefore, we hypothesized that transcriptional rewiring of the proteostasis network reflects an adaptive mechanism to preserve proteostasis in HSCs during chronic aging-associated stress.

To test this, we investigated whether specific proteostasis stress-related transcriptional programs are engaged during aging. Previous studies, which we confirmed (Fig. 1E,F), reported increased nuclear localization of Heat shock factor 1 (Hsf1) – a transcriptional regulator of many heat shock protein (HSP) genes (*115–117*) – in aged HSCs (*118*), though its transcriptional, molecular and functional impacts remained uncharacterized in this context. GSEA revealed robust enrichment of HSP gene expression in aged HSCs (Fig. 1G; p < 0.001). Since Hsf1 does not regulate all HSPs (*119–121*), we refined the list of 78 HSP genes to 57 that were previously validated as Hsf1 targets in at least some contexts (*119*) (table S3), and found that they were also significantly enriched in old HSCs (Fig. 1H; p < 0.001).

Since Hsf1 transcriptional programs can vary in a cell-type- and context-specific manner (*109, 119, 122–127*), we sought to define an HSC aging-specific Hsf1-HSP signature. To do this, we identified 50 HSP genes with increased expression (based on their positive rank metric score) in old versus young HSCs (fig. S1F, table S3). We then conditionally deleted *Hsf1* in hematopoietic cells by treating 6-week-old *Mx1*-*Cre^+^*;*Hsf1*^fl/fl^ (*Hsf1*^-/-^) mice (*128*) with polyinosinic:polycytidylic acid (pIpC), aged them to 24 months, and performed RNA-sequencing on HSCs. Of the 50 age-upregulated HSP genes, 29 genes had positive rank metric scores in aged WT compared to *Hsf1*^-/-^ HSCs (fig. S1G,H), and 22 of these were previously established Hsf1 targets (fig. S1I). Using this rigorous cutoff, we defined these 22 genes as an “Aging HSC Hsf1” signature (Fig. 1I, table S3, fig. S1E).

We validated this transcriptional signature across three independent datasets. First, we performed RNA-sequencing on young 3-month-old *Hsf1*⁻^/^⁻ and control HSCs. GSEA showed no enrichment of the “Aging HSC Hsf1” signature (Fig. 1J) in young WT versus *Hsf1*⁻^/^⁻ HSCs, suggesting that Hsf1-dependent transcriptional activation of these HSP genes is context-specific to aging (consistent with its age-dependent activation). Second, the “Aging HSC Hsf1” signature was significantly enriched in young HSCs following 18-hour *ex vivo* culture (Fig. 1K, GSE179412 (*118*)). Cultured HSCs experience significant proteostasis stress that induces nuclear accumulation of Hsf1 (*118*). Third, we analyzed single-cell RNA-sequencing data from young (21–24 years) and old (68–75 years) human Lineage^−^CD34^+^CD38^−^CD90^+^CD45RA^−^ HSCs and found that the “Aging HSC Hsf1” signature was significantly enriched in aged human HSCs (Fig. 1L, GSE188889 (*129*)).

Together, these findings demonstrate that Hsf1 is activated in HSCs in response to proteostasis stress during aging, driving a conserved transcriptional program of HSP coding genes in mice and humans.

### Hsf1 preserves proteostasis and self-renewal in old HSCs

To examine whether Hsf1 modulates proteostasis during HSC aging, we quantified protein synthesis and the accumulation of unfolded and misfolded proteins in HSCs from young and old *Hsf1*^-/-^ and control mice (Fig. 2A-F). Proteostasis was largely unaffected in young *Hsf1*^-/-^ HSCs (Fig. 2A-C), consistent with its minimal activation. However, old *Hsf1*^-/-^ HSCs exhibited a significant accumulation of unfolded and misfolded protein compared to age-matched controls (Fig. 2E,F), indicating that Hsf1 is specifically required to preserve proteostasis in old HSCs.

**Figure 2.**
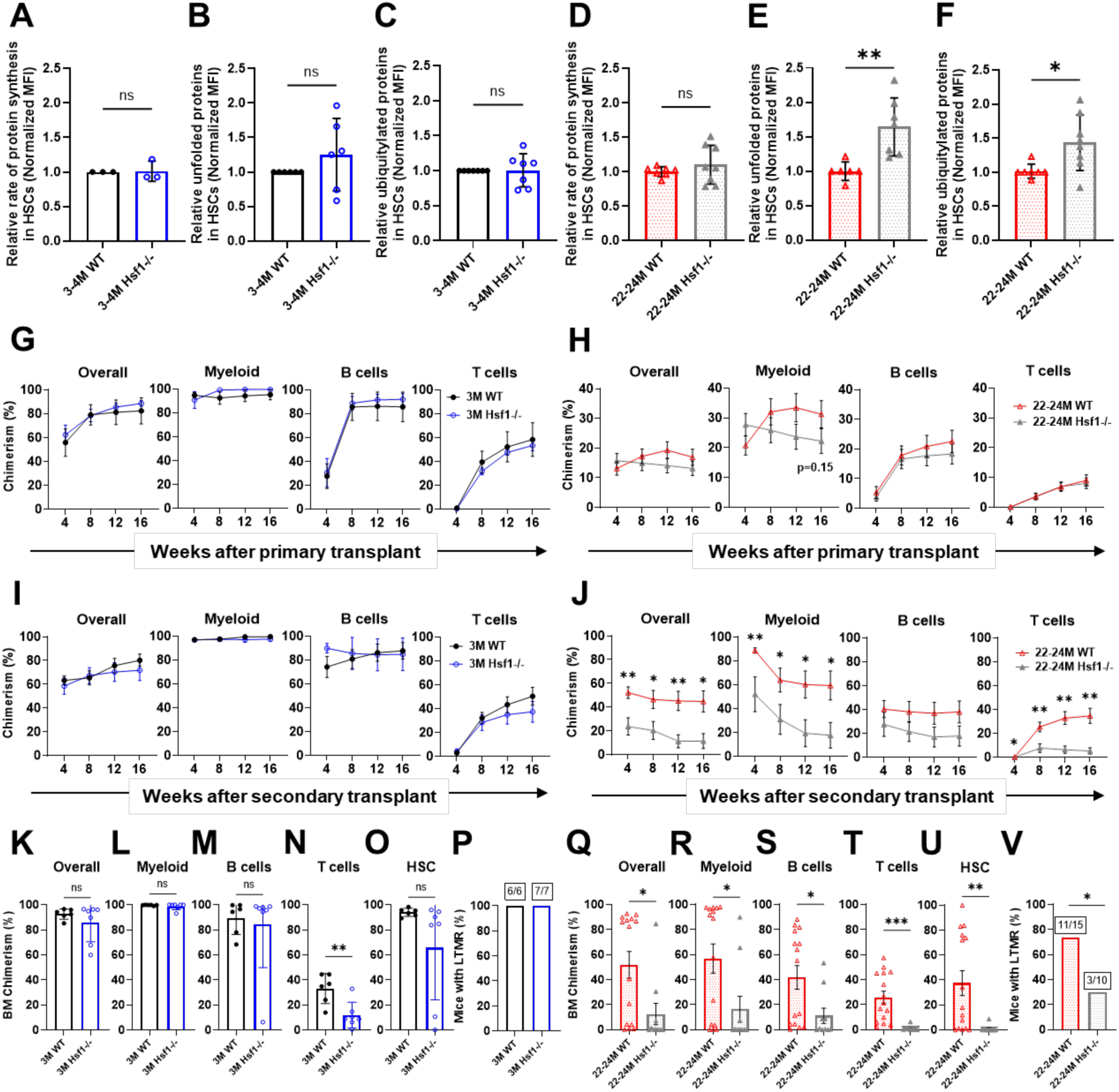
Hsf1 preserves proteostasis and self-renewal in old HSCs. **(A-F)** Relative (A,D) rates of protein synthesis (based on o-propargyl-puromycin incorporation), (B,E) unfolded protein abundance (based on tetraphenylethene maleimide fluorescence) and (C,F) misfolded protein abundance (based on polyubiquitylated protein) in (A-C) 3-4- and (D-F) 22-24-month-old WT and *Hsf1*^-/-^ HSCs (n=3-8 mice/age/genotype). **(G,H)** Donor hematopoietic, myeloid, B and T cell engraftment following transplantation of 25 HSCs from (G; n=2 donors, 4-5 recipients/genotype) 3- or (H; n=14-15 donors, 68-70 recipients/genotype) 22-24-month old WT and *Hsf1^-/-^* HSCs with 3×10^5^ recipient-type bone marrow cells into irradiated mice. **(I,J)** Donor hematopoietic, myeloid, B and T cell engraftment following secondary transplantation of 3×10^6^ bone marrow cells from recipients with median levels of reconstitution in (G) (I; n=2 donors, 6-7 recipients/genotype) or (H) (J; n=3 donors, 10-15 recipients/genotype). **(K-O)** Donor (K) total hematopoietic, (L) myeloid, (M) B cell, (N) T cell and (O) HSC engraftment in the bone marrow of secondary recipients from (I). **(P)** Frequency of secondary recipients from (I) that exhibited long-term multi-lineage reconstitution in peripheral blood. **(Q-U)** Donor (Q) total hematopoietic, (R) myeloid, (S) B cell, (T) T cell and (U) HSC engraftment in the bone marrow of secondary recipients from (J). **(V)** Frequency of secondary recipients from (J) that exhibited long-term multi-lineage reconstitution in peripheral blood. Data represent mean ± SD in (A-F, K-O, Q-U) and mean ± SEM in (G-J). Statistical significance was assessed using unpaired Student’s *t-*test in (A-O,Q-U) and Fisher’s exact test in (P,V). *p<0.05, **p<0.01, ***p<0.001

Given that proteostasis disruption can impair HSC function, we next tested whether Hsf1 is required for old HSC fitness and self-renewal. Steady state hematopoiesis remained largely normal in young (*118*) and old *Hsf1*^-/-^ mice (fig. S2A-Q). Competitive HSC transplantation assays (25 CD45.2^+^ HSCs vs. 3×10^5^ CD45.1^+^ bone marrow cells; fig. S2R) revealed no significant impact of *Hsf1*-deficiency on the long-term multilineage reconstituting activity of either young or old HSCs in primary recipients, although old *Hsf1*^-/-^ HSCs showed a trend toward reduced myeloid reconstitution (Fig. 2G,H; fig. S2S-V).

To directly assess self-renewal, we performed secondary transplants (fig. S2R). While *Hsf1*-deficiency had little effect on the serial reconstitution potential of young HSCs (Fig. 2I,K-P), old *Hsf1*^-/-^ HSCs exhibited a profound defect in long-term multilineage reconstitution in secondary recipients (Fig. 2J). This defect was characterized by a significant reduction in donor-derived cells in both the blood and bone marrow 16-weeks post-transplant, as well as a near-complete loss of donor-derived HSCs in the bone marrow (Fig. 2J, Q-U). Only 30% (3/10) of secondary recipients of old *Hsf1*^-/-^ HSCs exhibited long-term multilineage reconstitution compared to 73% (11/15) of recipients of age-matched controls (Fig. 2V; p < 0.05) and 100% of recipients of young *Hsf1*^-/-^ and control HSCs (Fig. 2P). Together, these data demonstrate that Hsf1 is essential to preserve both proteostasis and self-renewal capacity in old HSCs.

### Hsf1 promotes emergence of myeloid neoplasms by preserving proteostasis

Hsf1 has been implicated in the development and progression of multiple cancers (*123, 130–138*), including leukemias (*139–142*). While classically viewed as a stress response factor activated downstream of oncogenic signaling (*123, 130, 131, 143, 144*), our data raise the possibility that age-associated Hsf1 activation in HSCs might occur prior to disease onset to create a more permissive environment for malignant transformation.

To test whether oncogenic mutations drive Hsf1 activation, we assessed nuclear Hsf1 localization in HSCs from mice harboring early- (*Dnmt3a*^R878H^) (*6–8, 30*) and late-stage (*Nras*^G12D^) (*10–13, 145, 146*) AML-associated mutations. We used *Mx1*-*Cre^+^*;*Dnmt3a*^fl-R878H/+^ and *Mx1*-*Cre^+^*;*Nras*^fl-G12D/+^ mice (*147, 148*), as well as double mutants (*149, 150*) (*Mx1*-*Cre^+^*;*Dnmt3a*^fl-R878H/+^;*Nras*^fl-G12D/+^), in which mutations were induced by administering pIpC at 6 weeks of age. HSCs were analyzed at 6 months of age, prior to overt age-related Hsf1 activation (fig. S3A,B). Nuclear Hsf1 was not significantly increased in *Dnmt3a*^R878H^ HSCs, but was elevated in both *Nras*^G12D^ and *Dnmt3a*^R878H^;*Nras*^G12D^ HSCs (Fig. 3A,B). RNA-sequencing of *Dnmt3a*^R878H^;*Nras*^G12D^ and *Dnmt3a*^R878H^;*Nras*^G12D^;*Hsf1*^-/-^ HSCs (fig. S3C,D) revealed enrichment of HSP gene expression (fig. S3E) as well as the “Aging HSC Hsf1” signature in the presence of Hsf1 (Fig. 3C), confirming transcriptional activation and at least some transcriptional overlap with aging HSCs. In addition to HSP induction, *Dnmt3a*^R878H^;*Nras*^G12D^ HSCs exhibited transcriptional upregulation of multiple proteostasis pathways, which were diminished in the absence of *Hsf1* (Fig. 3D; table S4). Correspondingly, unfolded protein levels were significantly elevated in *Dnmt3a*^R878H^;*Nras*^G12D^;*Hsf1*^-/-^ HSCs (Fig. 3E), implicating Hsf1 in maintaining proteostasis under oncogenic stress.

**Figure 3.**
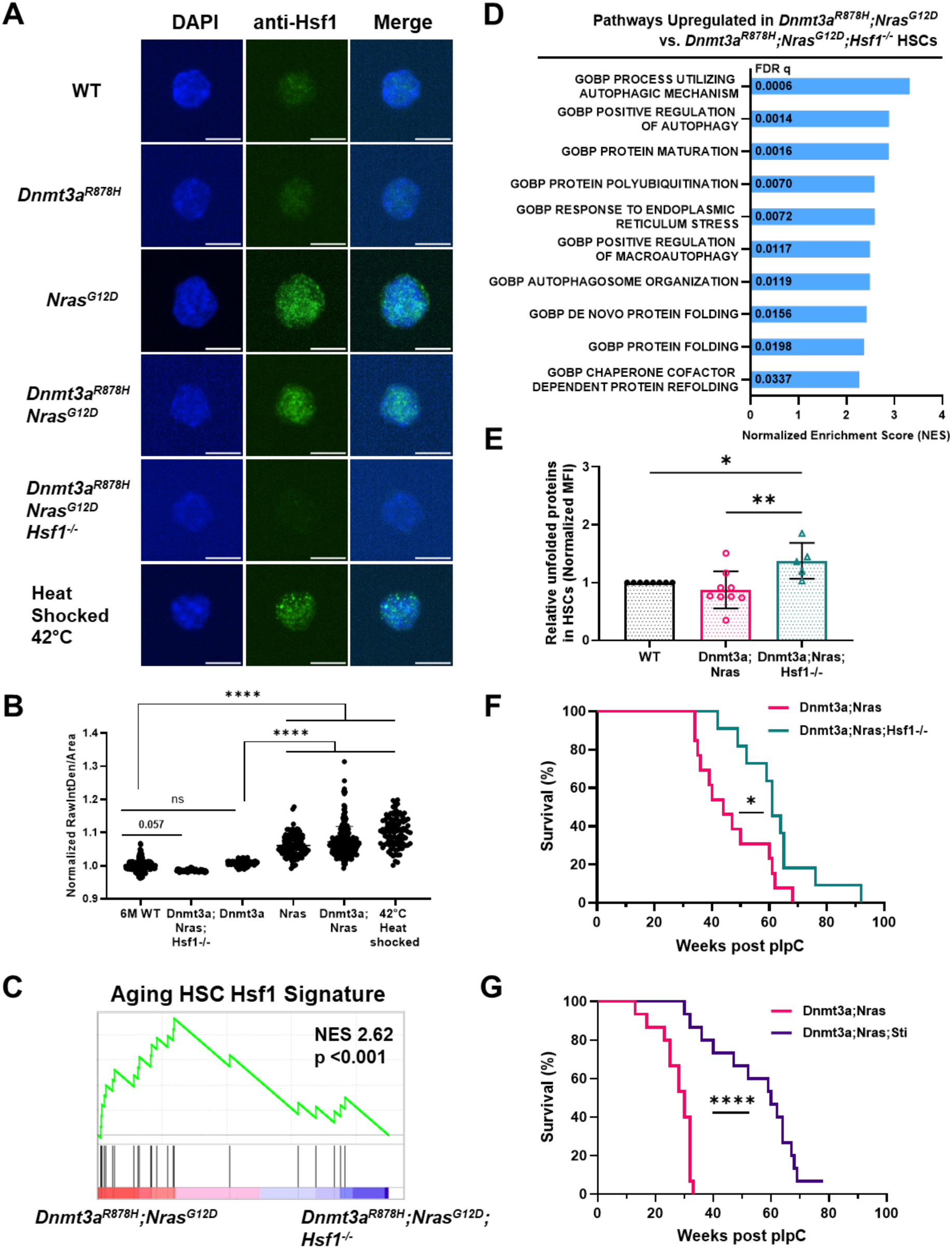
Hsf1 promotes emergence of myeloid neoplasms by preserving proteostasis. **(A,B)** Representative immunofluorescence images (A) and nuclear quantification (B) of Hsf1 in 6-month-old WT, *Dnmt3a^R878H^, Nras^G12D^, Dnmt3a^R878H^;Nras^G12D^,* and *Dnmt3a^R878H^;Nras^G12D^;Hsf1^-/-^*HSCs stained with DAPI and anti-Hsf1 (Alexa Fluor 488). WT CD150^−^CD48^−^LSK MPPs heat shocked at 42°C for >90 minutes were used as a positive control. Scale bars represent 5μm. Each dot represents an individual cell. **(C)** Gene set enrichment plot showing upregulation of “Aging HSC Hsf1” signature in *Dnmt3a^R878H^;Nras^G12D^* as compared to *Dnmt3a^R878H^;Nras^G12D^;Hsf1^-/-^* HSCs (n=3 mice/genotype). **(D)** GSEA from RNA-sequencing of 4-month-old *Dnmt3a^R878H^;Nras^G12D^*and *Dnmt3a^R878H^;Nras^G12D^;Hsf1^-/-^* HSCs. Select upregulated gene sets are shown with overlay of FDR q values (n=3 mice/genotype). **(E)** Relative unfolded protein abundance in WT, *Dnmt3a^R878H^;Nras^G12D^* and *Dnmt3a^R878H^;Nras^G12D^;Hsf1^-/-^* HSCs (n=5-9 mice per genotype). **(F,G)** Overall survival after pIpC treatment of chimeric mice engrafted with (F) *Mx1*-*Cre^+^*;*Dnmt3a*^fl-R878H/+^;*Nras*^fl-G12D/+^ or *Mx1*-*Cre^+^*;*Dnmt3a*^fl-R878H/+^;*Nras*^fl-G12D/+^;*Hsf1*^fl/fl^ or (G) *Mx1*-*_Cre_+*_;*Dnmt3a*_fl-R878H/+_;*Nras*_fl-G12D/+ _or *Mx1*-*Cre*_*+*_;*Dnmt3a*_fl-R878H/+_;*Nras*_fl-G12D/+_;*Aars*_sti/sti hematopoietic cells (n=11-15 mice/genotype). Data represent mean ± SD in (B,E). Statistical analysis was assessed using ordinary one-way ANOVA followed by Tukey-Kramer test for multiple comparisons in (B,E) and Log-rank (Mantel-Cox) test in (F,G). *p<0.05, **p<0.01, ****p<0.0001

To test whether Hsf1 promotes leukemogenesis in this context, we transplanted 10^7^ bone marrow cells from *Mx1*-*Cre^+^*;*Dnmt3a*^fl-R878H/+^;*Nras*^fl-G12D/+^ or *Mx1*-*Cre^+^*;*Dnmt3a*^fl-R878H/+^;*Nras*^fl-G12D/+^;*Hsf1*^fl/fl^ mice into irradiated recipients and treated with pIpC 4 weeks later. We confirmed robust (>95%) engraftment in all recipients (fig. S3F). Mice receiving *Dnmt3a*^R878H^;*Nras*^G12D^;*Hsf1*^-/-^ hematopoietic cells lived significantly longer than recipients of *Dnmt3a*^R878H^;*Nras*^G12D^ hematopoietic cells (Fig. 3F). Terminally ill mice of both genotypes displayed splenomegaly, with 75% (6/8) of spleens analyzed from *Dnmt3a^R878H^;Nras^G12D^*mice and 57% (4/7) of spleens analyzed from *Dnmt3a^R878H^;Nras^G12D^;Hsf1^-/-^*mice showing at least partial architectural effacement. Effaced spleens from both *Dnmt3a^R878H^;Nras^G12D^* and *Dnmt3a^R878H^;Nras^G12D^;Hsf1^-/-^*mice showed histomorphologic evidence of AML, although some pathological differences were observed. *Dnmt3a^R878H^;Nras^G12D^* mice showed predominance of immature myeloid or erythroid precursors (fig. S3G) while *Dnmt3a^R878H^;Nras^G12D^;Hsf1^-/-^* mice typically showed a predominance of immature erythroid precursors (fig. S3H). In addition, splenic megakaryocytes from *Dnmt3a^R878H^;Nras^G12D^*mice showed dysplasia (fig. S3G) while those from *Dnmt3a^R878H^;Nras^G12D^;Hsf1^-/-^* mice exhibited normal appearance (fig. S3H). These findings indicate that Hsf1 promotes the emergence and progression of fatal myeloid malignancies.

Since Hsf1 can promote cancer progression through pleiotropic mechanisms (*123*), it was unclear if proteostasis disruption contributed to impairing leukemogenesis in *Dnmt3a^R878H^;Nras^G12D^;Hsf1^-/-^*mice. Therefore, we tested whether proteostasis disruption itself was sufficient to impair leukemogenesis using an orthogonal model: *Aars*^sti/sti^ mice (*151*), which harbor a tRNA-editing defect that increases amino acid misincorporation and disrupts proteostasis without inducing significant Hsf1 activation. *Aars*^sti/sti^ HSCs accumulate unfolded proteins to a similar extent as *Dnmt3a*^R878H^;*Nras*^G12D^;*Hsf1*^-/-^ HSCs (fig. S3I) and do not upregulate the “Aging HSC Hsf1” transcriptional signature (fig. S3J, GSE141008 (*101*)). We transplanted 10^7^ bone marrow cells from *Mx1*-*Cre^+^*;*Dnmt3a*^fl-R878H/+^;*Nras*^fl-G12D/+^ or *Mx1*-*Cre^+^*;*Dnmt3a*^fl-R878H/+^;*Nras*^fl-G12D/+^;*Aars*^sti/sti^ mice into irradiated recipients and treated with pIpC 4 weeks later. Recipients of *Dnmt3a*^R878H^;*Nras*^G12D^;*Aars*^sti/sti^ cells had significantly prolonged survival (Fig. 3G), but ultimately developed terminal disease resembling that seen in *Dnmt3a*^R878H^;*Nras*^G12D^ mice (fig. S3K,L).

Together, these data reveal that some oncogenic mutations such as *Nras*^G12D^ can activate Hsf1, and that Hsf1 promotes leukemogenesis at least in part by preserving proteostasis in transformed HSCs. These data indicate that proteostasis mechanisms affecting normal stem cell fitness can similarly impair transformed HSCs.

### Proteostasis disruption impairs expansion of malignant progenitors

To further investigate how proteostasis disruption constrains leukemia development, we analyzed hematopoietic phenotypes in the context of the *Aars*^sti/sti^ mutation and *Hsf1*-deficiency in *Dnmt3a*^R878H^;*Nras*^G12D^ mice. As expected, *Dnmt3a*^R878H^;*Nras*^G12D^ mice exhibited significant expansion of multipotent and myeloid-restricted progenitors in the bone marrow (fig. S4A-N) and spleen, along with increased numbers of splenic HSCs (Fig. 4A-N). Notably, *both Aars*^sti/sti^ and *Hsf1*-deficiency markedly attenuated the expansion of several of these progenitor populations across hematopoietic compartments, though some effects did not reach the threshold of statistical significance (Fig. 4A-N; fig. S4A-N). These findings indicate that proteostasis maintenance is essential for expansion of malignant progenitors in this model.

**Figure 4.**
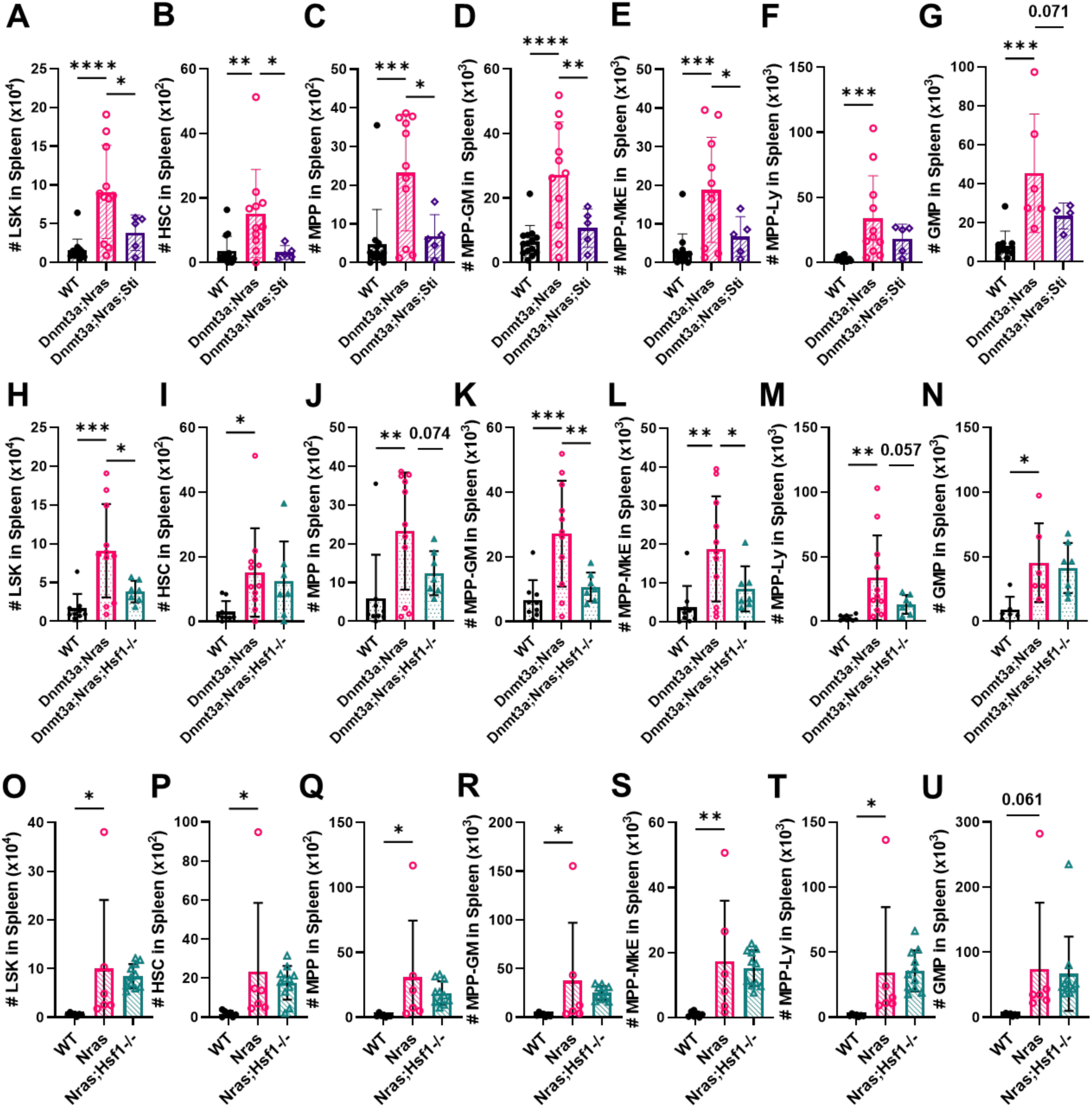
Proteostasis disruption impairs expansion of malignant progenitors in the spleen. **(A-G)** Number of (A) LSK cells, (B) CD150^+^CD48^−^LSK HSCs, (C) CD150^−^CD48^−^LSK MPPs, (D) CD150^−^CD48^+^LSK MPP^G/M^, (E) CD150^+^CD48^+^LSK MPP^Mk/E^, (F) CD135^+^LSK MPP^Ly^ and **(G)** CD34^+^CD16/32^high^CD127^−^Lineage^−^Sca1^−^cKit^+^ GMPs in the spleen of WT, *Dnmt3a*^R878H^;*Nras*^G12D^ and *Dnmt3a*^R878H^;*Nras*^G12D^;*Aars*^sti/sti^ mice (n=5-14 mice/genotype). **(H-N)** Number of (H) LSK cells, (I) HSCs, (J) MPPs, (K) MPP^G/M^, (L) MPP^Mk/E^, (M) MPP^Ly^ and (n) GMPs in the spleen of WT, *Dnmt3a*^R878H^;*Nras*^G12D^ and *Dnmt3a*^R878H^;*Nras*^G12D^;*Hsf1*^-/-^ mice (n=7-11 mice/genotype). **(O-U)** Number of (O) LSK cells, (P) HSCs, (Q) MPPs, (R) MPP^G/M^, (S) MPP^Mk/E^, (T) MPP^Ly^ and (U) GMPs in the spleen of WT, *Nras*^G12D^ and *Nras*^G12D^;*Hsf1*^-/-^ mice (n=6-11 mice/genotype). Data represent mean ± SD. Statistical analysis was assessed using ordinary one-way ANOVA with Fisher’s LSD relative to *Dnmt3a^R878H^;Nras^G12D^*in (A-N) or *Nras^G12D^* in (O-U). *p<0.05, **p<0.01, ***p<0.001, ****p<0.0001

Although the combination of *Dnmt3a*^R878H^ and *Nras*^G12D^ drive activation of Hsf1 and development of myeloid neoplasms that can progress to AML, *Nras*^G12D^ on its own is sufficient to activate Hsf1 (Fig. 3A,B) and drive hematopoietic progenitor expansion in *Mx1*-*Cre^+^*;*Nras*^fl-G12D/+^ mice (Fig. 4O-U; fig. S4O-U). However, in contrast to *Dnmt3a*^R878H^;*Nras*^G12D^ mice, *Hsf1*-deficiency did not substantially rescue hematopoietic stem and progenitor cell numbers in *Nras*^G12D^ mice (Fig. 4O-U; fig. S4O-U). These data indicate that while *Nras*^G12D^ can independently activate Hsf1 and drive progenitor expansion, the functional consequences of *Hsf1* loss require the presence of *Dnmt3a*^R878H^.

### Age-related Hsf1 activation promotes emergence of clonal hematopoiesis

Although the *Dnmt3a*^R878H^ mutation does not robustly activate Hsf1 on its own (Fig. 3A,B), our data suggest that Hsf1 activation can modulate the functional consequences of *Dnmt3a*^R878H^ in hematopoietic stem and progenitor cells. Since human *DNMT3A* mutations are among the earliest known events in leukemogenesis and are frequently detected in pre-malignant conditions such as age-related clonal hematopoiesis (*28, 30, 44, 45, 47, 48, 152*), we hypothesized that age-associated Hsf1 activation might promote the clonal expansion of *Dnmt3a* mutant HSCs.

To test this, we developed a murine model of age-related clonal hematopoiesis. We transplanted 10^7^ bone marrow cells from 3-month-old *Fgd5*^ZsGr-CreERT2+^;*Dnmta*^fl-R878H/+^ mice (CD45.2) into lethally irradiated, age-matched congenic WT CD45.1 recipients (Fig. 5A). Transplantation to establish chimeric recipients was necessary to circumvent the severe ulcerative dermatitis that develops in *Dnmta*^fl-R878H/+^ mice that precludes longitudinal aging studies. Four weeks post-transplant, recipients were pulsed with tamoxifen to induce stochastic recombination of the *Dnmt3a*^R878H^ allele in rare HSCs. Clonal expansion was tracked by quantifying the frequency of *Dnmt3a*^R878H^ HSCs in cohorts of mice sacrificed 1, 6, and 12-15 months post-tamoxifen, corresponding to 5-, 11-, and 17-20-month-old HSCs, respectively. Recombination was confirmed by genotyping colonies derived from single HSCs cultured in methylcellulose (Fig. 5A,B).

**Figure 5.**
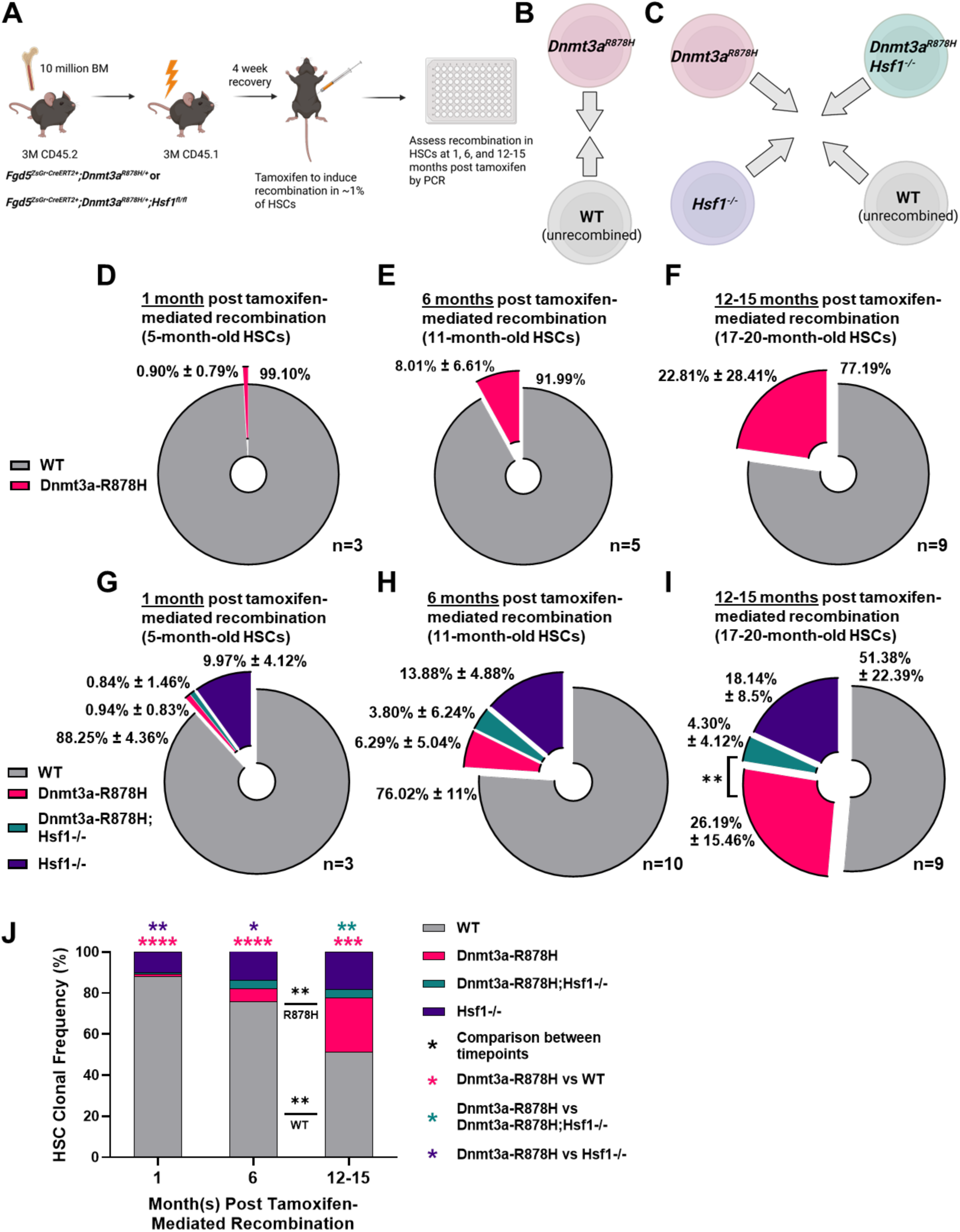
Age-related Hsf1 activation in HSCs promotes the emergence of *Dnmt3a^R878H^*-mediated clonal hematopoiesis. **(A)** Schematic of experimental clonal hematopoiesis setup establishing chimeric mice with hematopoietic cells transplanted from *Fgd5^ZsGr•CreERT2+^;Dnmt3a ^fl-R878H/+^* or *_Fgd5_ZsGr•CreERT2+_;Dnmt3a_ fl-R878H/+_;Hsf1_fl/fl* _mice._ **(B)** Schematic of clonal competition within *Fgd5^ZsGr•CreERT2+^;Dnmt3a^fl-R878H/+^* chimeras after tamoxifen administration. **(C)** Schematic of clonal competition within *Fgd5^ZsGr•CreERT2+^;Dnmt3a ^fl-R878H/+^;Hsf1^fl/fl^* chimeras after tamoxifen administration. **(D-F)** Clonal frequency of WT and *Dnmt3a^R878H^* HSCs at (D; n=3 mice) 1-, (E; n=5 mice) 6- and (F; n=9 mice) 12-15 months post tamoxifen-mediated recombination. **(G-I)** Clonal frequency of WT, *Dnmt3a^R878H^*, *Dnmt3a^R878H^;Hsf1^-/-^* and *Hsf1^-/-^* HSCs at (G; n=3 mice) 1-, (H; n=10 mice) 6- and (I; n=9 mice) 12-15-months post tamoxifen-mediated recombination. **(J)** Side-by-side comparison of HSC clonal composition by age from (G-I). Data represent mean ± SD in (D-I) and mean in (J). Statistical significance was assessed using paired Student’s *t-*test in (D-I), unpaired Student’s *t-*test in (J) for comparison between timepoints and ordinary one-way ANOVA with Fisher’s LSD relative to *Dnmt3a^R878H^*in (J) for comparison within a timepoint. *p<0.05, **p<0.01, ***p<0.001, ****p<0.0001

One month post-tamoxifen (corresponding to 5-month-old HSCs), *Dnmt3a*^R878H^ HSCs comprised 0.9% of the HSC pool (Fig. 5D). At 6 months post-tamoxifen (11-month-old-HSCs), *Dnmt3a*^R878H^ HSCs expanded to ∼8% of the HSC pool (Fig. 5E). At 12-15 months post-tamoxifen (17-20-month-old HSCs), *Dnmt3a*^R878H^ HSCs expanded further to ∼23% of the total HSC compartment (Fig. 5F; fig. S5A). This model recapitulates the dynamics of human clonal hematopoiesis, where rare mutations acquired early in life progressively expand to dominate the HSC pool with age (*23*).

To assess the role of Hsf1 in the clonal expansion process, we generated a parallel cohort of chimeric mice using donor cells from *Fgd5*^ZsGr-CreERT2+^;*Dnmta*^fl-R878H/+^;*Hsf1*^fl/fl^ mice. Following tamoxifen treatment, four distinct HSC populations were generated via stochastic recombination: **1) *Dnmt3a*^R878H^ only** (*Dnmt3a*^R878H^;*Hsf1*^fl/fl^), **2) *Hsf1*^-/-^ only** (*Dnmt3a*^fl-R878H/+^;*Hsf1*^-/-^), **3) Double mutant *Dnmt3a*^R878H^;*Hsf1*^-/-^**, and **4) WT** (unrecombined) (Fig. 5C). One month post-tamoxifen (5-month-old HSCs), the HSC compartment was comprised of 0.94% *Dnmt3a*^R878H^, 0.84% *Dnmt3a*^R878H^;*Hsf1*^-/-^, 9.97% *Hsf1*^-/-^ (suggesting higher recombination efficiency at the *Hsf1* locus), and 88.25% WT HSCs (Fig. 5G). At 6 months post-tamoxifen (11-month-old-HSCs), *Dnmt3a*^R878H^ clones expanded to 6.3% of the HSC pool, while *Dnmt3a*^R878H^;*Hsf1*^-/-^ HSCs exhibited modest expansion to 3.8% (Fig. 5H). This timepoint coincides with the approximate onset of robust Hsf1 activation in aging HSCs (*118*). By 12-15 months post-tamoxifen (17-20-month-old HSCs), *Dnmt3a*^R878H^ HSCs expanded to >26% of the HSC pool, whereas the double mutant *Dnmt3a*^R878H^;*Hsf1*^-/-^ HSCs failed to significantly expand, remaining at just 4.3% of the HSC compartment (Fig. 5I,J). These data indicate that age-related Hsf1 activation is required for the clonal expansion of *Dnmt3a*^R878H^ mutant HSCs and identify Hsf1 as a critical mediator of age-related clonal hematopoiesis.

### Hsf1 protects *Dnmt3a*^R878H^ HSCs from proteostasis and inflammatory stresses

Given Hsf1’s role in preserving proteostasis within aging and *Dnmt3a*^R878H^;*Nras*^G12D^ HSCs, we hypothesized that it might similarly promote clonal fitness of *Dnmta3a*^R878H^ HSCs by limiting the accumulation of unfolded and misfolded proteins. However, we found no significant differences in unfolded or misfolded protein abundance among WT, *Dnmt3a*^R878H^, and *Dnmt3a*^R878H^;*Hsf1*^-/-^ HSCs (Fig. 6A,B). An important limitation of this experiment is that *Dnmta*^fl-R878H/+^ mice develop ulcerative dermatitis that precludes their long-term aging, and so these analyses were performed in 4-month-old mice before maximal Hsf1 activation.

**Figure 6.**
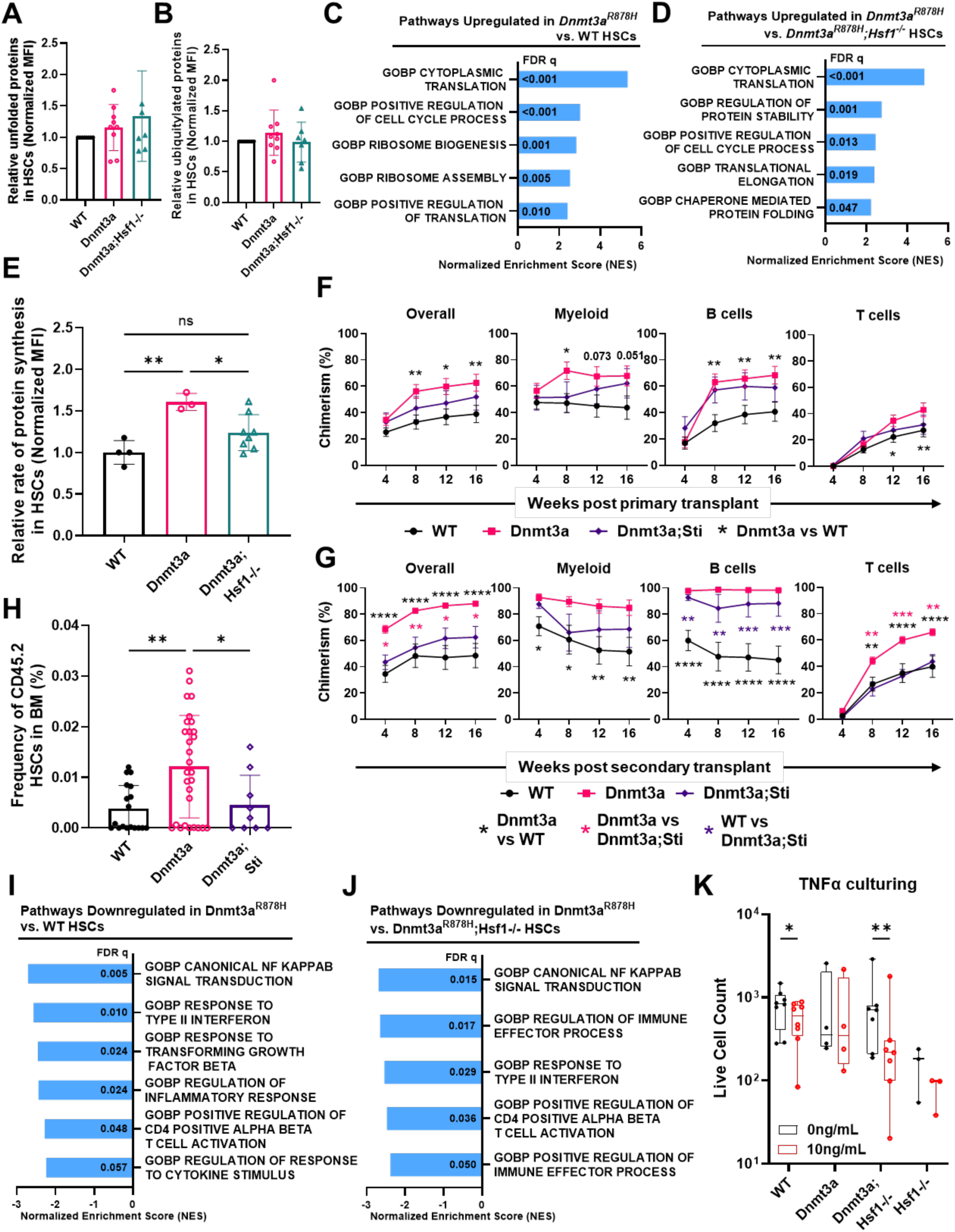
Hsf1 protects *Dnmt3a*^R878H^ HSCs from proteostasis and inflammatory stresses. **(A,B)** Relative (A) unfolded (based on tetraphenylethene maleimide fluorescence) and (B) misfolded protein abundance (based on polyubiquitylated protein) in WT, *Dnmt3a^R878H^*and *Dnmt3a^R878H^;Hsf1^-/-^* HSCs (n=7-9 mice/genotype). **(C,D)** Selected upregulated proteostasis-related pathways based on GSEA from RNA-sequencing in *Dnmt3a^R878H^* HSCs as compared to (C) WT and (D) *Dnmt3a^R878H^;Hsf1^-/-^*HSCs (n=2-3 mice/genotype). **(E)** Relative rates of protein synthesis (based on o-propargyl-puromycin incorporation) in WT, *Dnmt3a^R878H^* and *Dnmt3a^R878H^;Hsf1^-/-^* HSCs (n=3-8/genotype). **(F)** Donor hematopoietic, myeloid, B and T cell engraftment following transplantation of 15 HSCs from 4-month-old WT, *Dnmt3a^R878H^, Dnmt3a^R878H^;Aars^sti/sti^* HSCs with 3×10^5^ recipient-type bone marrow cells into irradiated mice (n=2-7 donors, 12-23 recipients/genotype). **(G)** Donor hematopoietic, myeloid, B and T cell engraftment following secondary transplantation of 3×10^6^ bone marrow cells from recipients with median levels of reconstitution in (F) (n=2-7 donors, 9-29 recipients/genotype). **(H)** Frequency of donor-derived HSCs in the bone marrow of secondary recipients from (G). **(I,J)** Selected downregulated inflammation-associated pathways based on GSEA from RNA-sequencing in *Dnmt3a^R878H^* HSC as compared to (I) WT and (J) *Dnmt3a^R878H^;Hsf1^-/-^* HSCs (n=2-3 mice/genotype). **(K)** Live cell numbers of WT, *Dnmt3a^R878H^, Dnmt3a^R878H^;Hsf1^-/-^* and *Hsf1^-/-^* HSCs after 7 days of culture with or without 10ng/mL TNFα. Each dot represents the mean of 2-4 technical replicates from one mouse (n=3-8 mice/genotype). Data represent mean ± SD in (A,B,E,H), mean ± SEM in (F,G) and median and quartiles in (K). Statistical analysis was assessed using an ordinary one-way ANOVA followed by Tukey-Kramer test for multiple comparisons in (A,B,E), two-way ANOVA (simple effects within rows) followed by Tukey-Kramer test for multiple comparisons in (F,G), ordinary one-way ANOVA followed by Dunnett’s test relative to *Dnmt3a^R878H^* in (H) and paired Student’s *t-*test in (K). *p<0.05, **p<0.01, ***p<0.001, ****p<0.0001

Despite this limitation, we sought to gain more mechanistic insight by performing RNA-sequencing on HSCs from 4-month-old mice of each genotype (fig. S6A-C). GSEA revealed significant upregulation of protein synthesis pathways in *Dnmt3a*^R878H^ HSCs relative to both WT and *Dnmt3a*^R878H^;*Hsf1*^-/-^ counterparts (Fig. 6C,D; tables S5,S6). Consistently, *Dnmt3a*^R878H^ HSCs exhibited significantly elevated protein synthesis rates, which were normalized in the absence of *Hsf1* (Fig. 6E). Increased protein synthesis at this level, such as in *Pten*^-/-^ HSCs, is known to impair HSC self-renewal by increasing the burden of unfolded/misfolded proteins (*100, 101*). Thus, our data suggest that Hsf1 enables *Dnmt3a*^R878H^ HSCs to sustain elevated protein synthesis (Fig. 6E) without compromising proteostasis (Fig. 6A,B). In the absence of *Hsf1*, *Dnmt3a*^R878H^ HSCs appear to adapt by suppressing protein synthesis back to near-WT levels to maintain proteostasis. *Dnmt3a*^R878H^;*Hsf1*^-/-^ HSCs also exhibit transcriptional upregulation of protein degradation and other proteostasis pathways that could contribute to proteostasis preservation in the absence of *Hsf1* (fig. S6D; table S8). These findings demonstrate that Hsf1 rebalances proteostasis through context-specific regulation of distinct proteostasis pathways.

To directly test whether proteostasis is required for the enhanced HSC self-renewal conferred by *Dnmt3a*^R878H^ (*153, 154*), we performed competitive serial transplantation (15 HSCs vs 3×10^5^ bone marrow cells) from 3-month-old *Mx1*-*Cre^+^*;*Dnmt3a*^fl-R878H/+^;*Aars*^sti/sti^, *Mx1*-*Cre^+^*;*Dnmt3a*^fl-R878H/+^, and WT control HSCs (pIpC treated at 6 weeks-of-age; fig. S6E). As expected, *Dnmt3a*^R878H^ HSCs gave elevated long-term reconstitution in both primary and secondary recipients (Fig. 6F,G). However, the *Aars*^sti/sti^ mutation significantly reduced the serial reconstituting activity of *Dnmt3a*^R878H^ HSCs back to near WT levels, although the effects were somewhat uneven across distinct lineages (Fig. 6F,G). The effect was most pronounced in secondary recipients and was accompanied by a significant reduction in donor-derived HSCs in the bone marrow, as there was a >3-fold increase in *Dnmt3a*^R878H^ HSCs in secondary recipients that was nearly completely abrogated in recipients of *Dnmt3a*^R878H^;*Aars*^sti/sti^ HSCs (Fig. 6H). These data indicate that disrupting proteostasis impairs the elevated self-renewal activity of *Dnmt3a*^R878H^ HSCs.

In addition to regulating protein synthesis pathways, RNA-sequencing revealed that inflammation-associated gene programs were markedly suppressed in *Dnmt3a*^R878H^ HSCs compared to both WT and *Dnmt3a*^R878H^;*Hsf1*^-/-^ HSCs (Fig. 6I,J; tables S7,S8). Given the importance of inflammatory resilience for clonal expansion *in vivo* (*155–160*), we hypothesized that Hsf1 might also confer protection for *Dnmt3a*^R878H^ HSCs against inflammatory stress. In vivo testing was not feasible since Hsf1 is largely activated during aging and the ulcerative dermatitis phenotype of *Dnmt3a*^fl-R878H/+^ mice precludes longitudinal aging studies in primary mice and is a confounding factor for studying inflammation. Instead, we cultured WT, *Dnmt3a*^R878H^, *Hsf1*^-/-^, and *Dnmt3a*^R878H^;*Hsf1*^-/-^ HSCs for 7 days in the presence or absence of the inflammatory cytokine TNFα (*157*). Culture induces Hsf1 activation in young adult HSCs and activates similar gene expression programs associated with Hsf1 activation during aging (Fig. 1K). As reported previously, TNFα reduced cell numbers in WT HSC cultures, but *Dnmt3a*^R878H^ HSCs were largely resistant to its effects (*157*) (Fig. 6K; fig. S6F). Notably, *Hsf1*-deficiency resensitized *Dnmt3a*^R878H^ HSCs to TNFα, resulting in significantly reduced cell numbers compared to vehicle-treated (PBS) *Dnmt3a*^R878H^;*Hsf1*^-/-^ controls (Fig. 6K; fig. S6F). These results suggest that Hsf1 protects *Dnmt3a*-mutant HSCs from proteostasis stress and confers resistance to inflammatory cues that would otherwise limit their clonal expansion.

Together, these findings demonstrate that Hsf1 activation is not merely a downstream response to oncogenic stress, but a crucial aging-associated event that establishes a more permissive proteostasis and inflammatory landscape for the emergence of clonal hematopoiesis. By buffering against both proteotoxic and inflammatory challenges, Hsf1 enables persistence and expansion of pre-leukemic clones carrying *Dnmt3a* mutations.

## Discussion

Functional declines in HSCs have been linked to nearly all molecular hallmarks of aging, with one notable exception: loss of proteostasis. This is surprising, given that proteostasis is fundamentally and preferentially important for preserving stem cell identity and self-renewal (*99, 101, 103*). Here, we uncover that proteostasis is not fully disrupted in aging HSCs, but rather selectively preserved through transcriptional reprogramming. We identify a critical role for Hsf1 and the heat shock response in maintaining proteome integrity in aging HSCs, protecting against unfolded and misfolded protein accumulation, and sustaining self-renewal potential.

In contrast to other tissues such as brain (*161*), liver (*162, 163*) and heart (*164*), where aging is associated with diminished Hsf1 activity and proteostasis collapse (*165–172*), HSCs exhibit increased Hsf1 activation with age (*118*). These findings reveal a striking cell-type- and context-specific divergence in proteostasis regulation. Whereas Hsf1 dysfunction drives aging in many tissues, its age-associated activation in HSCs represents a key adaptive mechanism that promotes stem cell fitness and longevity.

Despite this adaptation, Hsf1 activation alone is insufficient to fully prevent age-related HSC decline. Aging HSCs still exhibit myeloid-biased differentiation and reduced regenerative potential, likely due to concurrent aging hallmarks such as epigenetic dysregulation, impaired autophagy, and mitochondrial dysfunction (*51, 52, 54–58, 62, 65–67, 82, 103, 105, 106*).

Although Hsf1 can confer broad stress resistance, its endogenous activation is not sufficient to buffer the full complexity of aging-related perturbations. However, whether further enhancement of Hsf1 activation could confer additional fitness remains an area of future investigation.

In cancer, Hsf1 is frequently co-opted to drive malignant transformation, promoting stress tolerance, metastasis, and therapy resistance (*123, 131*). Many tumors display non-oncogene addiction to Hsf1 (*123, 143, 144*). Canonically, Hsf1 activation was viewed as a response to oncogenic stress *after* transformation (*123, 131, 143, 144*). We find that oncogenic Nras signaling alone is sufficient to induce Hsf1 activation in HSCs, although Hsf1 is not required for aberrant hematopoietic progenitor expansion in this context. In contrast, in *Dnmt3a*^R878H^;*Nras*^G12D^ double-mutant HSCs, *Hsf1*-deficiency results in proteotoxic stress, impairs clonal expansion, significantly delays the onset of malignancy, and prolongs survival in vivo.

Notably, *Dnmt3a* mutation alone does not robustly activate Hsf1. *DNMT3A* mutations, which are the most common drivers of age-related clonal hematopoiesis in people (*28, 30, 44, 45, 47, 48*), can arise early in life but only achieve significant clonal expansion in later decades (*23, 31*). Our findings suggest that physiological, age-associated Hsf1 activation provides a permissive environment that promotes the competitive advantage of *Dnmt3a*-mutant HSCs. Thus, cellular adaptations that preserve HSC integrity during aging are subverted by pre-leukemic clones to fuel clonal outgrowth and set the stage for leukemias to develop later in life by creating a more permissive state for cancer initiation and progression. While aging is a collapse of stress defense, cancer is its corruption.

Beyond its role in proteostasis, Hsf1 promotes stress resistance in *Dnmt3a*-mutant HSCs by conferring resilience to inflammatory stimuli, which is critical for clonal dominance during aging (*155–160*). Similar inflammatory resistance has also been described for other clonal hematopoiesis-associated mutations such as *TET2* (*155, 158, 173–176*), raising the possibility that Hsf1’s functions may extend across diverse pre-malignant contexts. Whether this represents a general mechanism of Hsf1 dependence in clonal hematopoiesis remains to be explored.

The pleiotropic nature of Hsf1 raises broader questions about its impact on stem cell biology and transformation. Recent studies have shown that *Dnmt3a*-mutant HSCs undergo metabolic reprogramming to rely more heavily on mitochondrial respiration (*177–179*). We also found that *Dnmt3a*-mutant HSCs upregulate gene programs associated with oxidative phosphorylation as compared to *Dnmt3a^R878H^;Hsf1^-/-^* HSCs (table S6), suggesting Hsf1 contributes to metabolic rewiring. This observation aligns with reports of Hsf1 influencing mitochondrial metabolism in AML and other cancers (*123, 140*), hinting at a potential link between proteostasis, metabolism, and clonal evolution that merits future investigation.

Collectively, our findings position Hsf1 not merely as a stress response factor, but as a central modulator of stem cell fate during aging and transformation. Hsf1 preserves HSC fitness under physiological stress, yet enables malignant progression when co-opted by oncogenic lesions. Hsf1 is not just a stress response factor, it is a culprit of aging and an architect of cellular resistance in pre-malignant stem cells and cancer.

Importantly, our data suggest that while Hsf1 supports proteostasis and self-renewal in aging HSCs, it is largely dispensable for steady-state hematopoiesis in older animals, indicating a possible therapeutic window. However, whether *Hsf1*-deficiency can be tolerated during hematopoietic stress, and how its inhibition would impact immune function or aging in other tissues, remain critical questions. An ideal therapeutic approach could be to identify and alleviate the intrinsic underlying stressors that drive Hsf1 activation in aging stem cells. By neutralizing these stressors, we may eliminate the selective pressure that drives Hsf1 activation and favors malignant clones, preserving healthy hematopoiesis and reducing the risk of transformation.

Such strategies could reshape the trajectory of aging and disease. By protecting the stem cell pool from underlying proteostasis stress, we could prevent the seeds of malignancy from ever taking root.

## Acknowledgements

J.A.M. is a scholar of the Leukemia and Lymphoma Society. The LJI Flow Cytometry Core, LJI Sequencing Core Facility, UCSD IGM Genomics Center, and UCSD Moores Cancer Center flow cytometry facility are supported by the NIH Shared Instrumentation Grant Program (S10 RR027366, S10 OD025052, S10 OD026929, S10 OD032316). The SCRM Flow Core is part of the UCSD Human Embryonic Stem Cell Core Facility and is supported by a CIRM Major Facilities grant (FA1-00607). We would like to thank the LJI Bioinformatics core and the Stem Cell Genomics Core at the Sanford Consortium for Regenerative Medicine for providing bioinformatics, sequencing, and microscopy services, E. Bennett for advice, Y. Hong for tetraphenylethene maleimide, and E. Christians for *Hsf1*^fl^ mice. Some figures were generated using BioRender.

## Funding

National Institutes of Health/National Heart, Lung, and Blood Institute fellowship F31HL170531 (FJZ)

National Institutes of Health/National Cancer Institute training grant 2T32CA067754 (FJZ)

National Cancer Institute Onco-Aging Consortium grant U01CA267031 (RAJS, JAM)

National Institute of Diabetes and Digestive and Kidney Diseases grant R01DK116951 (RAJS)

National Institute of Diabetes and Digestive and Kidney Diseases grant R01DK124775 (RAJS)

National Institute on Aging grant R01AG088725 (RAJS)

Cancer Stem Cell Consortium supported by the American Cancer Society and the Lisa Dean Moseley Foundation CSCC-RSG-23-994830-01-CSCC (RAJS)

Curebound Targeted Grant (RAJS)

UC San Diego Sanford Stem Cell Institute (RAJS) Sanford Stem Cell Discovery Center (RAJS)

UCSD Moores Cancer Center grant from National Institutes of Health/National Cancer Institute P30CA023100 (RAJS)

## Author contributions

R.A.J.S. and J.A.M. conceived the project. F.J.Z, R.A.J.S. and J.A.M. designed experiments. F.J.Z and R.A.J.S. wrote the manuscript. H.Y.W. provided histopathology analysis. F.J.Z. performed and analyzed all other experiments. M.K.L., H.C.W., A.T., and X.C. provided technical support. W.Y. supported bioinformatics analysis. M.J.S. supported mouse breeding and laboratory operations.

## Data and materials availability

All RNA-sequencing data generated for this study has been deposited in the National Center for Biotechnology Information Gene Expression Omnibus (NCBI GEO) and will be publicly available before publication. All previously existing datasets used for this study can be accessed through NCBI GEO using the indicated GSE accession number within the text (GSE179412, GSE188889, GSE141008). All other data are available in the main text or supplementary materials. All other relevant materials are available from the corresponding author upon reasonable request.

