## Supplemental Material for "Proteostasis Stress Drives Stem Cell Aging, Clonal Hematopoiesis and Leukemia"

**The PDF file include s:**

Materials and Methods
Figs. S1 to S6
References (180-197; listed under main text references)

**Other Supplementary Materials for this manuscript include the following:**

Tables S1 to S8 (provided as a separate .xlsx file with individual tables as different tabs)

#### Materials and Methods

Mice

*Mx1-Cre* (JAX 003556) (*180*), *Hsf1*$^{fl}$ *(128)*, *Dnmt3a*$^{fl\text{-}R878H}$ (JAX 032289) (*147*), *Nras*$^{fl\text{-}G12D}$ (JAX 008304) (*148*), *Aars*$^{sti/sti}$ *(151)*, and *Fgd5*$^{ZsGr\bullet CreERT2}$ (JAX 027789) (*181*) mice have been described previously. Mice were all backcrossed for at least ten generations onto a C57BL/6 background. *Hsf1*$^{fl/fl}$ mice were maintained as homozygous genotypes while all other alleles were maintained as heterozygous genotypes. *Aars*$^{sti}$ mice were maintained as heterozygous genotypes (*Aars*$^{sti/+}$) for breeding purposes but only homozygous genotypes (*Aars*$^{sti/sti}$) were used for experiments. C57BL6/J (CD45.2, JAX 000664) mice were used throughout the study. C57BL6.SJL (CD45.1, JAX 002014) mice were used in transplantation experiments. Both male and female mice were used in all studies. Mice used in these studies were between 3 and 24 months of age, as indicated.

Expression of *Mx1-Cre* was induced by six intraperitoneal injections of 10μg polyinosinic-polycytidylic acid (pIpC; Cytiva 27473201) administered every other day, beginning at approximately 5-7 weeks of age. All *Mx1-Cre*$^{-}$ controls (WT), *Aars*$^{sti/sti}$, and *Mx1-Cre*$^{+}$ experimental mice were treated with pIpC. Expression of *Fgd5*$^{ZsGr\bullet CreERT2}$ was induced by five intraperitoneal injections of 100mg/kg tamoxifen (Sigma T5648) administered every day, beginning at 4 weeks post-transplantation. Tamoxifen was reconstituted at 10mg/mL with 10% ethanol (Sigma E7023) in corn oil (Sigma C8267) overnight at 37°C and stored at -20°C protected from light.

All mice were housed in the vivarium at the University of California San Diego Moores Cancer Center or Sanford Consortium for Regenerative Medicine in specific pathogen-free conditions. All protocols were approved by the University of California San Diego Institutional Animal Care and Use Committee.

Flow cytometry and cell isolation

Mouse bone marrow cells were isolated by flushing the long bones (femurs and tibias) or by crushing the long bones, vertebrae, and pelvic bones with a mortar and pestle in $Ca^{2+}$- and $Mg^{2+}$-free Hank's buffered salt solution (HBSS; Corning 21-022-CV) supplemented with 2% (v/v) heat-inactivated bovine serum (Gibco 26170043). Spleens were prepared by crushing between frosted slides and resuspended in HBSS with 2% heat-inactivated bovine serum. All cells were filtered through a 40μm cell strainer to obtain single cell suspensions. Cell number and viability were assessed with a hemocytometer or TC20 Automated Cell Counter (BioRad) based on trypan blue exclusion. Cardiac blood was isolated immediately after euthanasia for complete blood count analysis using an Element HT5 (Heska).

For flow cytometric analysis and isolation of specific hematopoietic populations, cells were incubated with combinations of antibodies to the following cell-surface markers, conjugated to FITC, PE, PerCP-Cy5.5, APC, APC-eFluor 780, PE-Cy7, Alexa Fluor 700, or biotin (antibody clones are given in brackets in the following list): CD3 (17A2), CD4 (GK1.5), CD5 (53-7.3), CD8α (53-6.7), CD11b (M1/70), CD16/32 (FcγRII/III; 93), CD34 (RAM34), CD41 (MWReg30), CD43 (R2/60), CD45.1 (A20), CD45.2 (104), CD45R (B220; RA3-6B2), CD48 (HM48-1), CD71 (R17217), CD117 (c-Kit; 2B8), CD127 (IL7Rα; A7R34), CD135 (A2F10), CD150 (TC15-12F12.2), Ter119 (TER-119), Sca-1 (D7, E13-161.7), Gr-1 (RB6-8C5), and IgM

(II/41). Biotinylated antibodies were visualized by incubation with PE-Cy7 conjugated streptavidin (BioLegend 405206). All reagents were acquired from eBiosciences (Thermo Fisher Scientific) or BioLegend. All incubations were for 30-60 minutes on ice.

For isolation and analysis of mouse HSC and MPP populations, Lineage markers included CD3, CD5, CD8α, B220, Gr-1, and Ter119. HSCs were defined as CD150$^{+}$CD48$^{-}$Lineage$^{-}$Sca1$^{+}$cKit$^{+}$ (*110*), MPP as CD150$^{-}$CD48$^{-}$Lineage$^{-}$Sca1$^{+}$cKit$^{+}$ (*182, 183*), MPP$^{G/M}$ as CD150$^{-}$CD48$^{+}$Lineage$^{-}$Sca1$^{+}$cKit$^{+}$ (*183, 184*), MPP$^{Mk/E}$ as CD150$^{+}$CD48$^{+}$Lineage$^{-}$Sca1$^{+}$cKit$^{+}$ (*183, 184*) and MPP$^{Ly}$ as CD135$^{+}$Lineage$^{-}$Sca1$^{+}$cKit$^{+}$ (*183, 184*). For analysis of mouse CMPs, GMPs, and MEPs, Lineage markers included CD3, CD4, CD8α, CD11b, B220, Gr-1, and Ter119. CMPs were defined as CD34$^{+}$CD16/32$^{low}$CD127$^{-}$Lineage$^{-}$Sca1$^{-}$cKit$^{+}$ (*185*), GMPs as CD34$^{+}$CD16/32$^{high}$CD127$^{-}$Lineage$^{-}$Sca1$^{-}$cKit$^{+}$ (*185*) and MEPs as CD34$^{-}$CD16/32$^{low}$CD127$^{-}$Lineage$^{-}$Sca1$^{-}$cKit$^{+}$ (*185*). For lymphoid lineages, IgM$^{+}$ B cells were defined as IgM$^{+}$B220$^{+}$ (*186*), Pre-B cells as IgM$^{-}$CD43$^{-}$B220$^{+}$ (*186*), Pro-B cells as IgM$^{-}$CD43$^{+}$B220$^{+}$ (*186*) and T cells as CD4$^{+}$ or CD8$^{+}$. For myeloid lineages, erythrocytes were defined as Ter119$^{+}$, myeloid cells (GM) as Gr1$^{+}$CD11b$^{+}$ and megakaryocytes (Mk) as CD41$^{+}$.

Mouse HSCs were pre-enriched by selecting c-Kit$^{+}$ cells using paramagnetic microbeads and an autoMACS magnetic separator (Miltenyi Biotec) before sorting. Non-viable cells were excluded from sorts and analyses using 1μg/mL 4',6-diamidino-2-phenylindole (DAPI). Analysis of protein synthesis rates (*100, 187*), ubiquitylated protein (pan-ubiquitin antibody, clone FK2, Sigma 04263) (*103*), and unfolded protein (tetraphenylethene maleimide) (*103, 113*) by flow cytometry were performed exactly as previously described.

Cell sorting was performed on a BD FACSAriaII or BD FACSAria Fusion. Sorted fractions were double sorted (yield then purity precision) or sorted with single cell precision to ensure high purity. Data acquisition for analysis was performed using BD LSRII, BD LSRFortessa X-20 with HTS or BD FACSymphony A1. Data were analyzed using FlowJo (BD) software.

<u>Transplantation assays</u>

Adult recipient mice (CD45.1) were administered a minimum lethal dose of radiation using a Mark I Cesium source irradiator (J.L. Sheperd) or X-Rad320 X-ray irradiator (Precision X-Ray) to deliver two doses of 475-550 rad (950-1,100 rad in total) 4 hours apart. Cells were injected into the retro-orbital venous sinus of anesthetized recipients. Transplanted mice were administered drinking water with Baytril (250mg/L; Bayer 08711170) for the first 4 weeks post-transplantation.

For primary HSC transplants, 15 or 25 freshly isolated CD150$^{+}$CD48$^{-}$Lineage$^{-}$Sca-1$^{+}$c-Kit$^{+}$ HSCs (CD45.2) and 3x10$^{5}$ CD45.1 bone marrow cells were transplanted into lethally irradiated recipients. Blood was obtained from the tail veins of recipient mice every 4 weeks for 16 weeks after transplantation. Red blood cells were lysed with ammonium chloride potassium buffer. The remaining cells were stained with antibodies against CD45.2, CD45.1, CD45R (B220), CD11b, CD3, and Gr-1 to assess donor-cell engraftment (chimerism). After sixteen weeks post-transplantation, bone marrow of primary recipients was analyzed for HSC engraftment and multilineage chimerism.

For secondary transplants, $3x10^6$ bone marrow cells collected from primary recipients were transplanted non-competitively into lethally irradiated recipient mice (CD45.1). Primary recipients used for secondary transplantation had long-term multilineage reconstitution by donor cells and median levels of donor-cell reconstitution for the treatments from which they originated. Mice were considered long-term multilineage reconstituted if they exhibited >0.5% donor-derived peripheral blood hematopoietic, B, T, and myeloid cells 16 weeks post-transplantation. Blood was obtained from the tail veins of recipient mice every 4 weeks for 16 weeks post-transplantation. After 16 weeks post-transplantation, bone marrow of secondary recipients was analyzed for HSC engraftment and multilineage chimerism.

For survival studies (Fig. 3F,G), $10^7$ bone marrow cells (CD45.2) were transplanted from *Mx1-Cre*$^{+}$*;Dnmt3a*$^{fl\text{-}R878H/+}$*;Nras*$^{fl\text{-}G12D/+}$, *Mx1-Cre*$^{+}$*;Dnmt3a*$^{fl\text{-}R878H/+}$*;Nras*$^{fl\text{-}G12D/+}$*;Hsf1*$^{fl/fl}$ or *Mx1-Cre*$^{+}$*;Dnmt3a*$^{fl\text{-}R878H/+}$*;Nras*$^{fl\text{-}G12D/+}$*;Aars*$^{sti/sti}$ mice into lethally irradiated recipients (CD45.1). Four weeks post-transplantation, expression of *Mx1-Cre* was induced by six intraperitoneal injections of 10µg pIpC administered every other day. Blood was obtained from the tail veins of recipient mice every 4 weeks until recipients were moribund (euthanized) or found dead.

For clonal competition studies (Fig. 6D-I), $10^7$ bone marrow cells (CD45.2) were transplanted from *Fgd5*$^{\text{ZsGr-CreERT2+}}$;*Dnmt3a*$^{fl\text{-}R878H/+}$ or *Fgd5*$^{\text{ZsGr-CreERT2+}}$;*Dnmt3a*$^{fl\text{-}R878H/+}$*;Hsf1*$^{fl/fl}$ mice into lethally irradiated recipients (CD45.1). Four weeks post-transplantation, expression of *Fgd5*$^{ZsGr\bullet CreERT2}$ was induced by five intraperitoneal injections of 100mg/kg tamoxifen administered every day.

Methylcellulose colony genotyping

Single freshly isolated $CD150^{+}CD48^{-}Lineage^{-}Sca\text{-}1^{+}c\text{-}Kit^{+}$*Fgd5*$^{ZsGr\bullet CreERT2+}$ HSCs from bone marrow of recipients in clonal competition studies were sorted per well of a 96-well plate containing methylcellulose culture medium (StemCell Technologies M3434) and incubated at 37°C in 5% $CO_2$ and air and constant humidity. PBS was added between wells to ensure the methylcellulose did not dry out. DNA was extracted from colonies 13-15 days after plating by first resuspending methylcellulose in PBS and then transferring to 1.5mL microcentrifuge tubes. Cells were washed twice with PBS in microcentrifuge tubes to remove methylcellulose and transferred back to a 96-well plate for DNA extraction in a thermocycler. DNA was extracted with DirectPCR Mouse Tail Lysis Reagent (Viagen Biotech 102-T) with 30µg Proteinase K (0.9 units, Sigma P6556) at 55°C for 4 hours and 85°C for 45 minutes. 1µL of lysate (DNA) was used for colony genotyping and a range of 66-167 colonies were genotyped per mouse.

Colonies were PCR genotyped using the following primers for *Dnmt3a*$^{fl\text{-}R878H/+}$ recombination (Common F, 5'-CTCCTTGGATTTGAGGAGGA-3'; WT R, 5'-TGCACATGAGAACTGGATGG-3'; Mut R, 5'-ATTAAGGGCCAGCTCATTCC-3') and *Hsf1*$^{fl/fl}$ recombination (F, 5'-CTAGTCAGTCCCTAGAGATGACCAG-3'; R, 5'-GTTGTGGTCAGCTCCTGTC-3').

Immunostaining and confocal microscopy

For analysis of Hsf1 activation, freshly isolated $CD150^{+}CD48^{-}Lineage^{-}Sca\text{-}1^{+}c\text{-}Kit^{+}$ HSCs were sorted and plated on VistaVision HistoBond Microscope Slides (VWR 16004-406) with RetroNectin (Clontech Labs/Takara Bio T100A). RetroNectin was diluted to 12.5µg/mL in

DPBS (Corning 21-030-CV) and 5μL was placed for each sample on top of microscope slides. To allow RetroNectin to bind, slides were incubated at 37°C for at least 30 minutes in a humid environment. Excess RetroNectin was removed before plating cells on top of the RetroNectin droplet to allow cells to bind to the microscope slide. RetroNectin slides with cells were incubated at 37°C for 15 minutes. Cells were then directly fixed in 4% paraformaldehyde in PBS (Electron Microscopy Sciences 157-4) for 15 minutes at room temperature. Cells were washed three times with PBS and permeabilized with PBS supplemented with 0.1% Triton X-100 (VWR 80503-490), 0.03% Tween20 (Sigma P9416) and 1% BSA (Sigma A9647) for 20 minutes at room temperature. Cells were washed three times with PBS supplemented with 0.03% Tween20. Cells were blocked with PBS supplemented with 1% BSA and 0.03% Tween20 for 60 minutes at room temperature. Samples were incubated with anti-Hsf1 antibody (Cell Signaling, Rabbit polyclonal 4356) at 1:500 dilution in blocking solution overnight at 4°C. Cells were washed three times with PBS supplemented with 0.03% Tween20 and incubated with Alexa Fluor 488 conjugated donkey anti-Rabbit secondary antibody (Thermo Fisher A-21206) at 1:500 dilution in blocking solution with 2μg/mL DAPI for 60 minutes at room temperature. Cells were washed three times with PBS and mounted in ProLong Diamond Antifade Mountant (Molecular Probes P36961) with No. 1.5 Gold Seal Cover Glass (Electron Microscopy Sciences 6379110). All processes were done on the RetroNectin microscope slides with a hydrophobic barrier (Newcomer Supply PAP Pen Liquid Blocker 6506).

Images were acquired using a DMi8 microscope (Leica) attached to a spinning disk Dragonfly 200 High Speed Confocal Microscope System (Andor/Oxford Instruments) with the Fusion software (Andor). All images were acquired using a 63X oil-immersion objective with Zyla cameras and excited using 405nm (DAPI) and 488nm (Alexa Fluor 488) lasers. Z-stack images were taken with the same confocal settings for Alexa Fluor 488 with minor adjustments for DAPI between samples. Raw images were processed with FIJI (ImageJ) and imported as a Hyperstack. Max Intensity ZProjection was generated for each Hyperstack. Blue (DAPI) and green (anti-Hsf1/Alexa Fluor 488) channels were merged and Alexa Fluor 488 intensity was quantified based on co-localization of DAPI and Hsf1. RawIntDen/Area was used for quantification and at least 100 cells were quantified per sample if possible. The same Brightness/Contrast settings were used across samples within an experiment for Alexa Fluor 488 with minor adjustments for DAPI between samples.

<u>Inflammation cultures</u>
100 $CD150^{+}CD48^{-}Lineage^{-}Sca\text{-}1^{+}c\text{-}Kit^{+}$ HSCs were sorted (single cell precision) into a 96-well plate containing 150μL of Ham's F-12 media (Gibco 11765054) supplemented with 1x penicillin-streptomycin (Corning 30002-CI), 10mmol/L HEPES (Gibco 15630080), 2mM L-Glutamine (Gibco 25030081), 1x insulin-transferrin-selenium-ethanolamine (Gibco 51500056), 100ng/mL recombinant murine TPO (Peprotech 315-141), 10ng/mL recombinant murine SCF (Peprotech 250-03), and 1mg/mL polyvinyl alcohol (Sigma 363146) as previously described (*157, 188*). HSCs from individual mice were sorted into 4-8 independent wells. Half the wells were treated with 10ng/mL recombinant murine TNFα (Peprotech 315-01A). The HSCs were cultured for 7 days at 37°C and 5% $CO_2$. Additional recombinant TNFα or PBS was spiked into cultures on days 4 and 6. On day 7, 100μL of Ham's F-12 with a final concentration of 1μg/mL DAPI was added into each well immediately before analysis by flow cytometry for live cell count.

Histopathology

Mouse tissues were fixed in 10% Buffered Formalin (Fisher Scientific 23-305510) for at least 24 hours. Tissues were then washed with MilliQ water and stored in 70% ethanol before embedding. Tissues were paraffin embedded, cut, and hematoxylin and eosin (H&E) stained according to standard protocols by the Moores Cancer Center Biorepository and Tissue Technology Core. 10 individual fields at 400X were scored per H&E slide using a light microscope.

RNA-sequencing

For RNA-sequencing analysis, total RNA was extracted from up to 5,000 HSCs using the RNeasy Plus Micro Kit (Qiagen 74034). Illumina mRNA libraries were prepared using the SMART-seq protocol (*189*) (Takara Bio). 2-2.6μL of total RNA was used in the SMART-seq protocol (100-400pg). For Figure 1 and Figure S1, 18 cycles of PCR were performed for the cDNA preamplification step and 12 cycles were performed for the tagmentation library preparation. The resulting libraries were pooled and deep sequenced in two lanes on the Illumina NovaSeq 6000 using paired-end reads with both forward and reverse read lengths of 50 nucleotides (50-115M reads per condition). For Figures 3 and 6 and Figures S3 and S6, 10 cycles of PCR were performed for the cDNA preamplification step and 18 cycles were performed for the tagmentation library preparation. The resulting libraries were double-sided size selected (0.6-1x), pooled and deep sequenced in one lane on the Illumina NovaSeq X Plus using paired-end reads with both forward and reverse read lengths of 75 nucleotides (80-107M reads per condition). The reads that passed Illumina filters were filtered for reads aligning to tRNA, rRNA, adapter sequences, and spike-in controls. The reads were then aligned to GRCm38 reference genome using STAR (v2.6.1c) (*190*). DUST scores were calculated with PRINSEQ Lite (v 0.20.3) (*191*) and low complexity reads (DUST > 4) were removed from the BAM files. The alignments results were parsed via SAMtools (*192*) to generate SAM files. Read counts to each genomic feature were obtained with htseq-count program (v0.7.1) (*193*) using the "union" option. After removing absent features (zero counts in all samples), the raw counts were then imported to Bioconductor package DESeq2 (v1.24.0) (*194*) to identify differentially expressed genes among samples. P-values for differential expression were calculated using the Wald test for differences between the base means of two conditions. These P-values were then adjusted for multiple test correction using Benjamini Hochberg algorithm. We considered genes differentially expressed between two groups of samples when the DESeq2 analysis resulted in an adjusted P-value of <0.05 and the difference in gene expression was at least 1.5-fold. GSEA (*112*) was done using the "GseaPreranked" method with "classic" scoring scheme. MSigDB gene sets (*195, 196*) were downloaded for mouse Gene Ontology (*197*) analysis. Rank files for each DESeq2 comparison were generated by assigning a rank of negative $\log_{10}$(pValue) to genes with log2FoldChange greater than zero and a rank of positive $\log_{10}$(pValue) to genes with a log2FoldChange less than zero. For analysis against the "Aging HSC Hsf1" signature gene set (table S3), "GseaPreranked" method with "classic" scoring scheme was also used.

Statistical Methods

Group data are represented by mean ± standard deviation (SD), except for transplantation data which are represented as mean ± standard error of the mean (SEM). To test statistical significance between two samples, two-tailed Student's *t*-tests were used. When multiple samples were compared to one another, statistical significance was assessed using repeated-measures one-way ANOVA followed by Tukey-Kramer or Dunnett's test for multiple comparisons or two-way ANOVA (simple effects within rows) followed by Tukey-Kramer test for multiple comparisons. Statistical significance comparing overall number of mice with long-term multilineage reconstitution was assessed using Fisher's exact test. Statistical significance comparing differences in survival was calculated using a Log-rank (Mantel-Cox) test. Statistical tests were performed using GraphPad Prism software. The specific type of test used for each figure panel is described in the figure legends, except for RNA sequencing data which is described in the "RNA-sequencing" method section. For normalized data, statistical tests were performed using $log_{10}$-transformed data to prevent data skewing. No randomization or blinding was used in any experiments. The only mice excluded from any experiment were those that died after transplantation. In the case of measurements in which variation tends to be low (for sample, HSC frequency), we generally examined 3-6 young (3-4 month-old) mice and ~10 old (22-24 month-old) or mutant ($Dnmt3a^{fl\text{-}R878H/+}$;$Nras^{fl\text{-}G12D/+}$) mice. In the case of measurements in which variation among experiments tends to be higher (for example, reconstitution assays), we examined larger numbers of mice (>10 per experiment). We performed multiple independent experiments on different days with multiple biological replicates to ensure reproducibility of our findings.

#### Figure S1

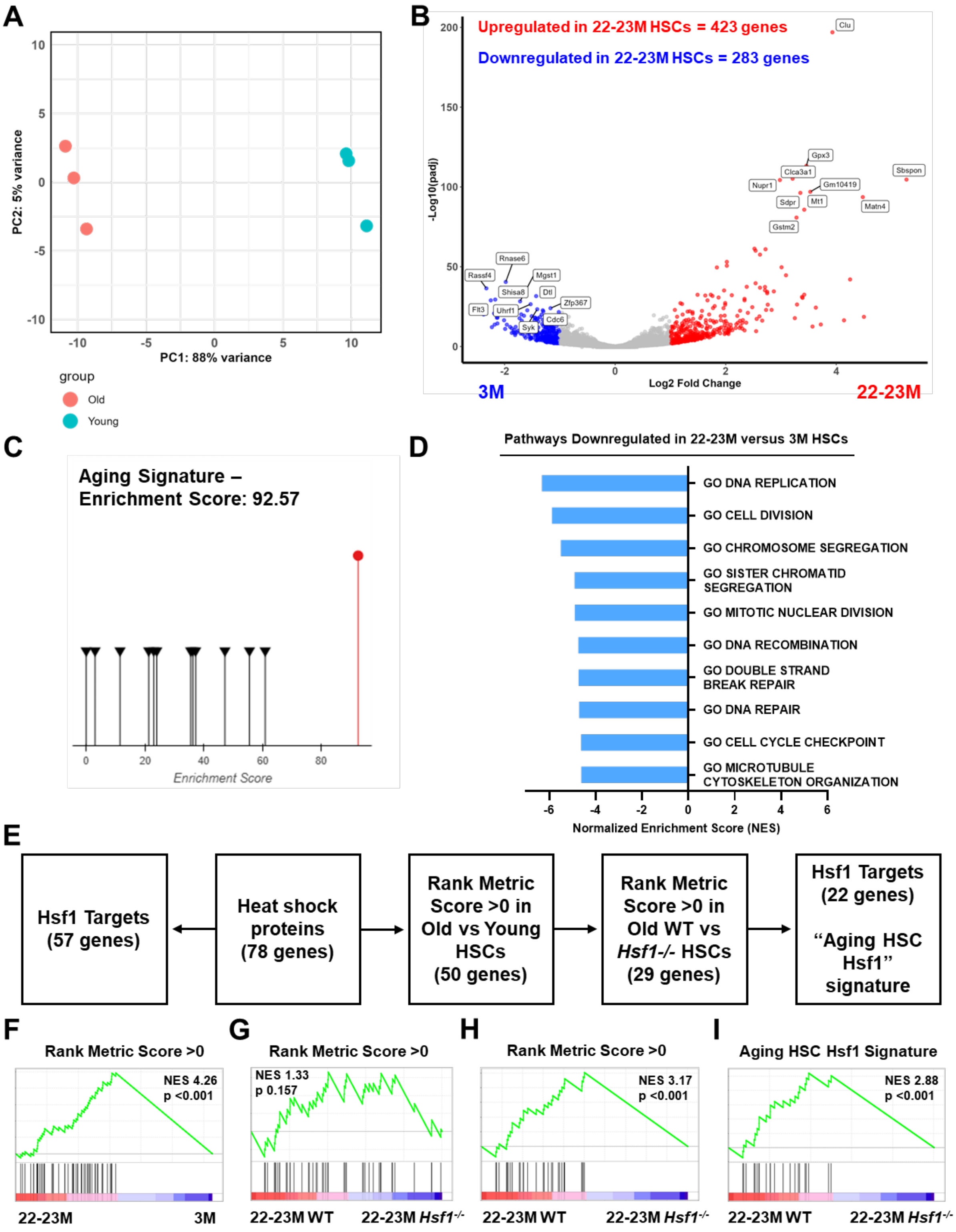
A
PC2: 5% variance
PC1: 88% variance
group
Old
Young
B
Upregulated in 22-23M HSCs = 423 genes
Downregulated in 22-23M HSCs = 283 genes
-Log10(padj)
Log2 Fold Change
3M
22-23M
Clu
Gpx3
Clca3a1
Nupr1
Gm10419
Sbspon
Sdpr
Mt1
Matn4
Gstm2
Rnase6
Rassf4
Mgst1
Shisa8
Dtl
Flt3
Uhrf1
Zfp367
Cdc6
Syk
C
Aging Signature –
Enrichment Score: 92.57
Enrichment Score
D
Pathways Downregulated in 22-23M versus 3M HSCs
GO DNA REPLICATION
GO CELL DIVISION
GO CHROMOSOME SEGREGATION
GO SISTER CHROMATID SEGREGATION
GO MITOTIC NUCLEAR DIVISION
GO DNA RECOMBINATION
GO DOUBLE STRAND BREAK REPAIR
GO DNA REPAIR
GO CELL CYCLE CHECKPOINT
GO MICROTUBULE CYTOSKELETON ORGANIZATION
Normalized Enrichment Score (NES)
E
Hsf1 Targets (57 genes)
Heat shock proteins (78 genes)
Rank Metric Score >0 in Old vs Young HSCs (50 genes)
Rank Metric Score >0 in Old WT vs Hsf1-/- HSCs (29 genes)
Hsf1 Targets (22 genes)
"Aging HSC Hsf1" signature
F
Rank Metric Score >0
NES 4.26
p <0.001
22-23M
3M
G
Rank Metric Score >0
NES 1.33
p 0.157
22-23M WT
22-23M Hsf1-/-
H
Rank Metric Score >0
NES 3.17
p <0.001
22-23M WT
22-23M Hsf1-/-
I
Aging HSC Hsf1 Signature
NES 2.88
p <0.001
22-23M WT
22-23M Hsf1-/-

**Figure S1 (Related to Figure 1). Transcriptional rewiring of aging HSCs.**
**(A)** PCA plot of 22-23- and 3-month-old HSCs (n=3 mice/age).
**(B)** Volcano plot showing upregulated and downregulated genes in 22-23- versus 3-month-old HSCs (n=3 mice/age). Genes highlighted in red and blue correspond to absolute fold change of 2 and $p_{adj} < 0.05$.
**(C)** Enrichment score comparing upregulated genes in 22-23-month-old HSCs (red line) compared to a database of an established mouse HSC aging signature (*111*) (black lines).
**(D)** GSEA in 3- and 22-23-month-old HSCs from RNA-sequencing. Selected downregulated gene sets are shown. False discovery rates (FDR) q values for all graphed gene sets are <0.001 (n=3 mice/age).
**(E)** Schematic showing the development of the "Aging HSC Hsf1" signature.
**(F)** Gene set enrichment plot showing upregulation of heat shock protein (HSP) genes with rank metric score >0 in 22-23- old HSCs versus 3M old HSCs from Fig. 1G (n=3 mice/age)
**(G)** Gene set enrichment plot showing HSP genes from (F) in 22-23-month-old WT and *Hsf1*$^{-/-}$ HSCs (n=3 mice/genotype).
**(H)** Gene set enrichment plot showing HSP genes with rank metric score >0 in 22-23M WT HSCs from (G) (n=3 mice/genotype).
**(I)** Gene set enrichment plot showing "Aging HSC Hsf1" signature in 22-23M WT HSCs versus *Hsf1*$^{-/-}$ HSCs (n=3 mice/genotype).

### Figure S2

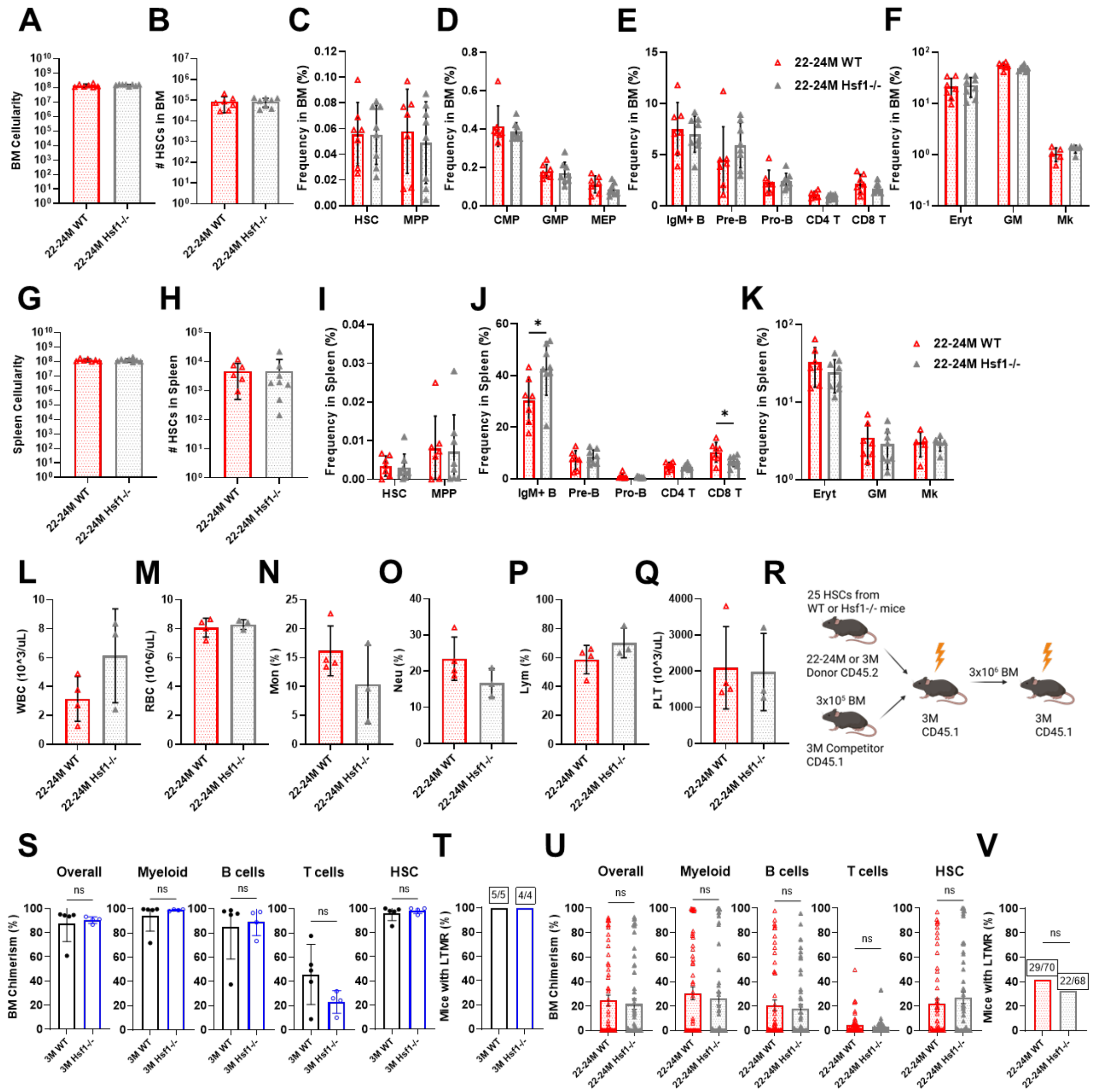

A
B
C
D
E
F
BM Cellularity
### HSCs in BM
Frequency in BM (%)
HSC
MPP
CMP
GMP
MEP
IgM+ B
Pre-B
Pro-B
CD4 T
CD8 T
Eryt
GM
Mk
22-24M WT
22-24M Hsf1-/-
G
H
I
J
K
Spleen Cellularity
### HSCs in Spleen
Frequency in Spleen (%)
L
M
N
O
P
Q
R
WBC (10^3/uL)
RBC (10^6/uL)
Mon (%)
Neu (%)
Lym (%)
PLT (10^3/uL)
25 HSCs from WT or Hsf1-/- mice
22-24M or 3M Donor CD45.2
3x10⁵ BM
3M Competitor CD45.1
3M CD45.1
3x10⁶ BM
S
T
U
V
Overall
Myeloid
B cells
T cells
HSC
ns
BM Chimerism (%)
Mice with LTMR (%)
3M WT
3M Hsf1-/-
5/5
4/4
29/70
22/68

**Figure S2 (Related to Figure 2). *Hsf1*-deficiency does not significantly impact steady state hematopoiesis during aging.**

**(A-K)** Hematopoietic analysis of 22-24-month-old WT and $Hsf1^{-/-}$ mice, including (A) bone marrow and (G) spleen cellularity, (B,H) HSC number in the bone marrow (2 femurs + 2 tibias) and spleen, frequency of (C,I) $CD150^{+}CD48^{–}Lineage^{–}Sca1^{+}cKit^{+}$ HSCs and $CD150^{–}CD48^{–}Lineage^{–}Sca1^{+}cKit^{+}$ MPPs, (D) $CD34^{+}CD16/32^{low}CD127^{–}Lineage^{–}Sca1^{–}cKit^{+}$ CMPs, $CD34^{+}CD16/32^{high}CD127^{–}Lineage^{–}Sca1^{–}cKit^{+}$ GMPs, $CD34^{–}CD16/32^{low}CD127^{–}Lineage^{–}Sca1^{–}cKit^{+}$ MEPS, (E,J) $IgM^{+}B220^{+}$ B cells, $IgM^{+}B220^{+}CD43^{-}$ pre-B, $IgM^{+}B220^{+}CD43^{+}$ pro-B cells, $CD4^{+}$ and $CD8^{+}$ T cells, (F,K) $Ter119^{+}$ erythroid (Ery), $Gr1^{+}CD11b^{+}$ myeloid (GM) and $CD41^{+}$ megakaryocyte (Mk) lineage cells in the bone marrow and spleen (n=3-8 mice/genotype).

**(L-Q)** Complete blood counts from 22-24-month-old WT and $Hsf1^{-/-}$ mice including (L) white blood cells, (M) red blood cells, (N) monocyte frequency, (O) neutrophil frequency, (P) lymphocyte frequency, and (Q) platelet number.

**(R)** Schematic for competitive, serial transplantation of 25 3- and 22-24-month-old WT or $Hsf1^{-/-}$ HSCs into irradiated recipients. Data are shown in Fig. 2G-V.

**(S)** Donor total hematopoietic, myeloid, B cell, T cell and HSC engraftment in the bone marrow of primary recipients from Fig. 2G.

**(T)** Frequency of primary recipients from Fig. 2G that exhibited long-term multi-lineage reconstitution in peripheral blood.

**(U)** Donor total hematopoietic, myeloid, B cell, T cell and HSC engraftment in the bone marrow of primary recipients from Fig. 2H.

**(V)** Frequency of primary recipients from Fig. 2H that exhibited long-term multi-lineage reconstitution in peripheral blood.

Data represent mean ± SD in (A-Q,S,U). Statistical significance was assessed using an unpaired Student's *t*-test in (A-Q,S,U) and Fisher's exact test in (T,V). $^{*}p<0.05$

**Figure S3**

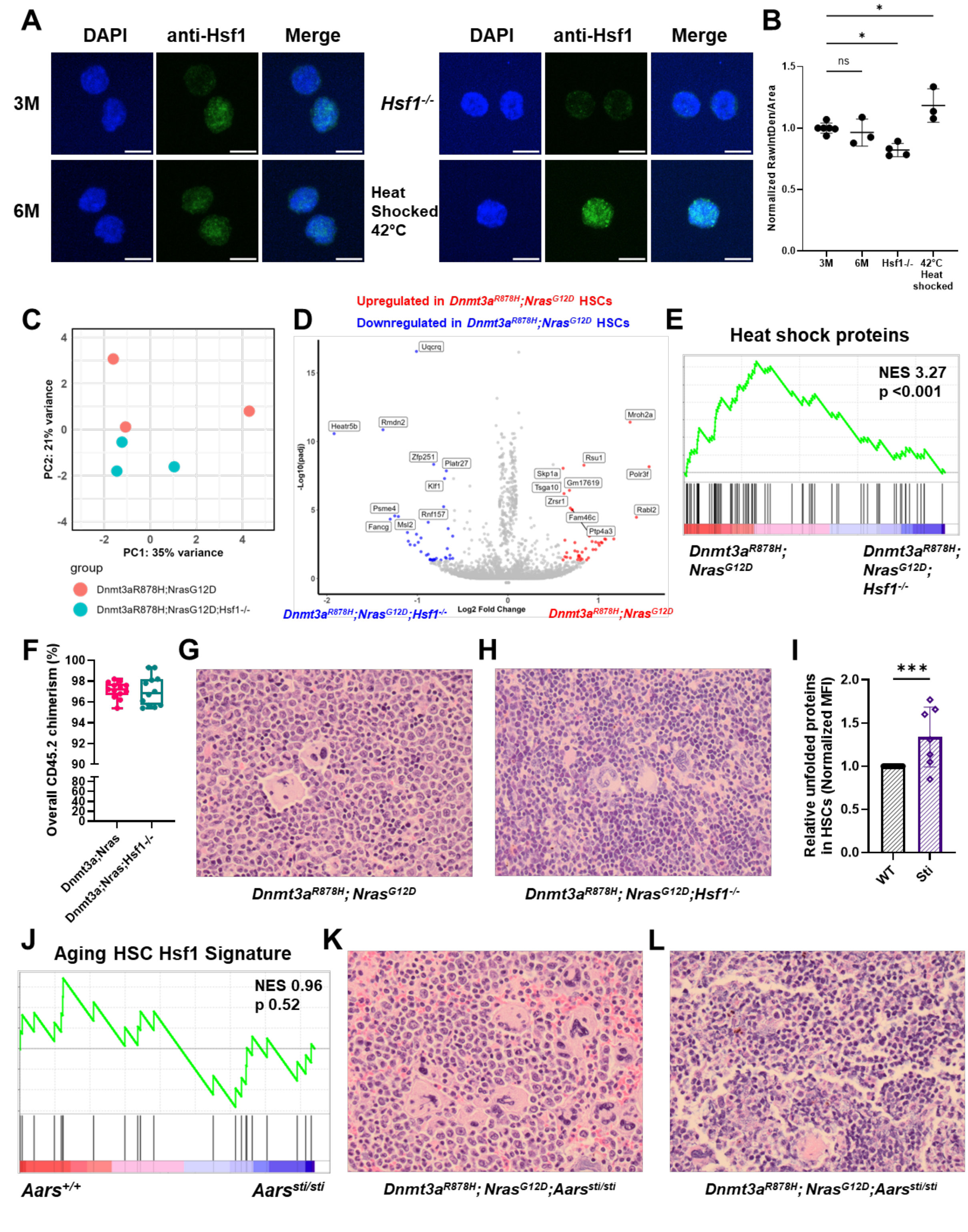
A
DAPI
anti-Hsf1
Merge
3M
6M
Hsf1-/-
Heat Shocked 42°C
B
Normalized RawIntDen/Area
ns
*
*
3M
6M
Hsf1-/-
42°C Heat shocked
C
PC1: 35% variance
PC2: 21% variance
group
Dnmt3aR878H;NrasG12D
Dnmt3aR878H;NrasG12D;Hsf1-/-
D
Upregulated in Dnmt3aR878H;NrasG12D HSCs
Downregulated in Dnmt3aR878H;NrasG12D HSCs
-Log10(padj)
Log2 Fold Change
Dnmt3aR878H;NrasG12D;Hsf1-/-
Dnmt3aR878H;NrasG12D
E
Heat shock proteins
NES 3.27
p <0.001
Dnmt3aR878H; NrasG12D
Dnmt3aR878H; NrasG12D; Hsf1-/-
F
Overall CD45.2 chimerism (%)
Dnmt3a;Nras
Dnmt3a;Nras;Hsf1-/-
G
H
Dnmt3aR878H; NrasG12D
Dnmt3aR878H; NrasG12D;Hsf1-/-
I
Relative unfolded proteins in HSCs (Normalized MFI)
***
WT
Sti
J
Aging HSC Hsf1 Signature
NES 0.96
p 0.52
Aars+/+
Aarssti/sti
K
L
Dnmt3aR878H; NrasG12D;Aarssti/sti
Dnmt3aR878H; NrasG12D;Aarssti/sti

**Figure S3 (Related to Figure 3). Impact of *Hsf1*-deficiency and proteostasis disruption on gene expression and pathology in *Dnmt3a*$^{R878H}$;*Nras*$^{G12D}$-driven neoplasms.**
**(A,B)** Representative immunofluorescence images (A) and normalized nuclear quantification (B) of Hsf1 in 3- and 6-month-old WT and *Hsf1*$^{-/-}$ HSCs stained with DAPI and anti-Hsf1 (Alexa Fluor 488). WT MPPs heat shocked at 42°C for >90 minutes were used as a positive control. Scale bars represent 5µm. (B) Each dot represents the average of ~100 cells per mouse normalized to 3-month-old HSCs.
**(C)** PCA plot from RNA-sequencing of 4-month-old *Dnmt3a*$^{R878H}$*;Nras*$^{G12D}$ and *Dnmt3a*$^{R878H}$*;Nras*$^{G12D}$*;Hsf1*$^{-/-}$ HSCs (n=3 mice/genotype).
**(D)** Volcano plot showing upregulated and downregulated genes in *Dnmt3a*$^{R878H}$*;Nras*$^{G12D}$ as compared to *Dnmt3a*$^{R878H}$*;Nras*$^{G12D}$;*Hsf1*$^{-/-}$ HSCs (n=3 mice/genotype). Genes highlighted in red and blue correspond to absolute fold change of 1.5 and $p_{adj} < 0.05$.
**(E)** Gene set enrichment plot showing upregulation of HSP genes in *Dnmt3a*$^{R878H}$*;Nras*$^{G12D}$ as compared to *Dnmt3a*$^{R878H}$*;Nras*$^{G12D}$;*Hsf1*$^{-/-}$ HSCs (n=3 mice/genotype).
**(F)** Donor hematopoietic cell engraftment in chimeric mice from survival studies in Fig. 3F grafted with *Mx1-Cre*$^{+}$;*Dnmt3a*$^{fl\text{-}R878H/+}$;*Nras*$^{fl\text{-}G12D/+}$ or *Mx1-Cre*$^{+}$;*Dnmt3a*$^{fl\text{-}R878H/+}$;*Nras*$^{fl\text{-}G12D/+}$;*Hsf1*$^{fl/fl}$ hematopoietic cells (n=12-13 mice/genotype).
**(G)** Representative image of spleen from *Dnmt3a*$^{R878H}$*;Nras*$^{G12D}$ moribund chimeras (from Fig. 3F) at 400X magnification. Shown is near total effacement of splenic architecture, which is replaced by immature granulocytic precursors with dysplastic megakaryocytes.
**(H)** Representative image of spleen from *Dnmt3a*$^{R878H}$*;Nras*$^{G12D}$*;Hsf1*$^{-/-}$ moribund chimeras (from Fig. 3F) at 400X magnification. Shown is effacement of splenic architecture, which is replaced by immature erythroid precursors with normal appearing megakaryocytes.
**(I)** Relative unfolded protein abundance in 4-month-old old WT and *Aars*$^{sti/sti}$ HSCs (n=7-15 mice/genotype).
**(J)** Gene set enrichment plot of "Aging HSC Hsf1" signature in young *Aars*$^{+/+}$ (WT) and *Aars*$^{sti/sti}$ HSCs (from GSE141008).
**(K,L)** Representative images of spleens from *Dnmt3a*$^{R878H}$*;Nras*$^{G12D}$*;Aars*$^{sti/sti}$ moribund chimeras (from Fig. 3G) at 400X magnification. Shown is near total (K) or total (L) effacement of splenic architecture, which is replaced by immature granulocytic precursors with dysplastic megakaryocytes (K) or immature erythroid precursors (L).
Data represent mean ± SD in (B,F,I). Statistical significance was assessed using ordinary one-way ANOVA followed by Dunnett's test for multiple comparisons relative to 3M in (B) and unpaired Student's *t*-test in (I). *p<0.05, ***p<0.001

**Figure S4**

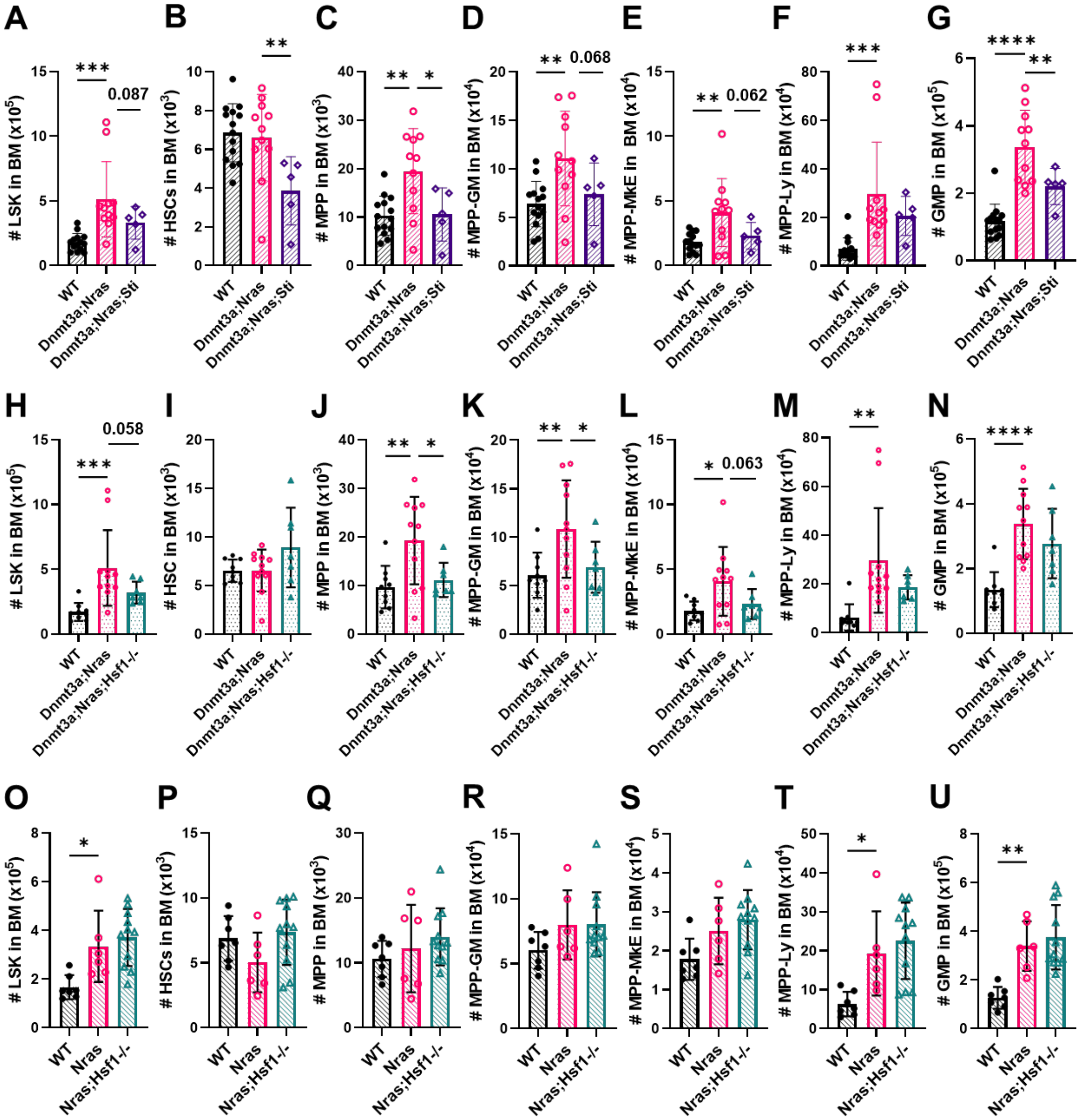
A
B
C
D
E
F
G
H
I
J
K
L
M
N
O
P
Q
R
S
T
U
WT
Dnmt3a;Nras
Dnmt3a;Nras;Sti
Dnmt3a;Nras;Hsf1-/-
Nras
Nras;Hsf1-/-
0.087
0.068
0.062
0.058
0.063

**Figure S4 (Related to Figure 4). Proteostasis disruption impairs expansion of malignant progenitors in the bone marrow.**

**(A-G)** Number of (A) $Lineage^{-}Sca^{+}c\text{-}Kit^{+}$ (LSK) cells, (B) $CD150^{+}CD48^{-}$LSK HSCs, (C) $CD150^{-}CD48^{-}$LSK MPPs, (D) $CD150^{-}CD48^{+}$LSK $MPP^{G/M}$, (E) $CD150^{+}CD48^{+}$LSK $MPP^{Mk/E}$, (F) $CD135^{+}$LSK $MPP^{Ly}$ and (G) $CD34^{+}CD16/32^{high}CD127^{-}Lineage^{-}Sca1^{-}cKit^{+}$ GMPs in the bone marrow of WT, *Dnmt3a*$^{R878H}$;*Nras*$^{G12D}$ and *Dnmt3a*$^{R878H}$;*Nras*$^{G12D}$;*Aars*$^{sti/sti}$ mice (n=5-14 mice/genotype).

**(H-N)** Number of (H) LSK cells, (I) HSCs, (J) MPPs, (K) $MPP^{G/M}$, (L) $MPP^{Mk/E}$, (M) $MPP^{Ly}$ and (N) GMPs in the bone marrow of WT, *Dnmt3a*$^{R878H}$;*Nras*$^{G12D}$ and *Dnmt3a*$^{R878H}$;*Nras*$^{G12D}$;*Hsf1*$^{-/-}$ mice (n=7-11 mice/genotype).

**(O-U)** Number of (O) LSK cells, (P) HSCs, (Q) MPPs, (R) $MPP^{G/M}$, (S) $MPP^{Mk/E}$, (T) $MPP^{Ly}$ and (U) GMPs in the bone marrow of WT, *Nras*$^{G12D}$ and *Nras*$^{G12D}$;*Hsf1*$^{-/-}$ mice (n=6-11 mice/genotype).

Data represent mean ± SD. Statistical analysis was assessed using ordinary one-way ANOVA with Fisher's LSD relative to *Dnmt3a*$^{R878H}$;*Nras*$^{G12D}$ in (A-N) or *Nras*$^{G12D}$ in (O-U). *$p<0.05$, **$p<0.01$, ***$p<0.001$, ****$p<0.0001$

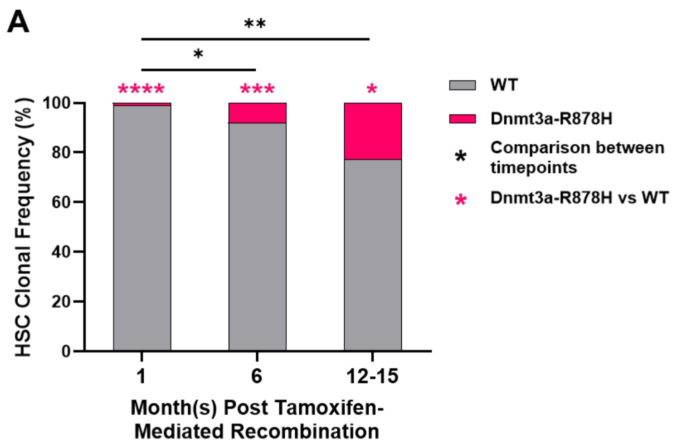


**Figure S5 (Related to Figure 5). Establishing a mouse model of *Dnmt3a*$^{R878H}$-mediated age-related clonal hematopoiesis.**

**(A)** Side-by-side comparison of HSC clonal composition by age from Fig. 5D-F.

Data represent mean. Statistical significance was assessed using paired Student's *t*-test within a timepoint and Mann-Whitney test for comparison between timepoints. *p<0.05, **p<0.01 ***p<0.001, ****p<0.0001

### Figure S6

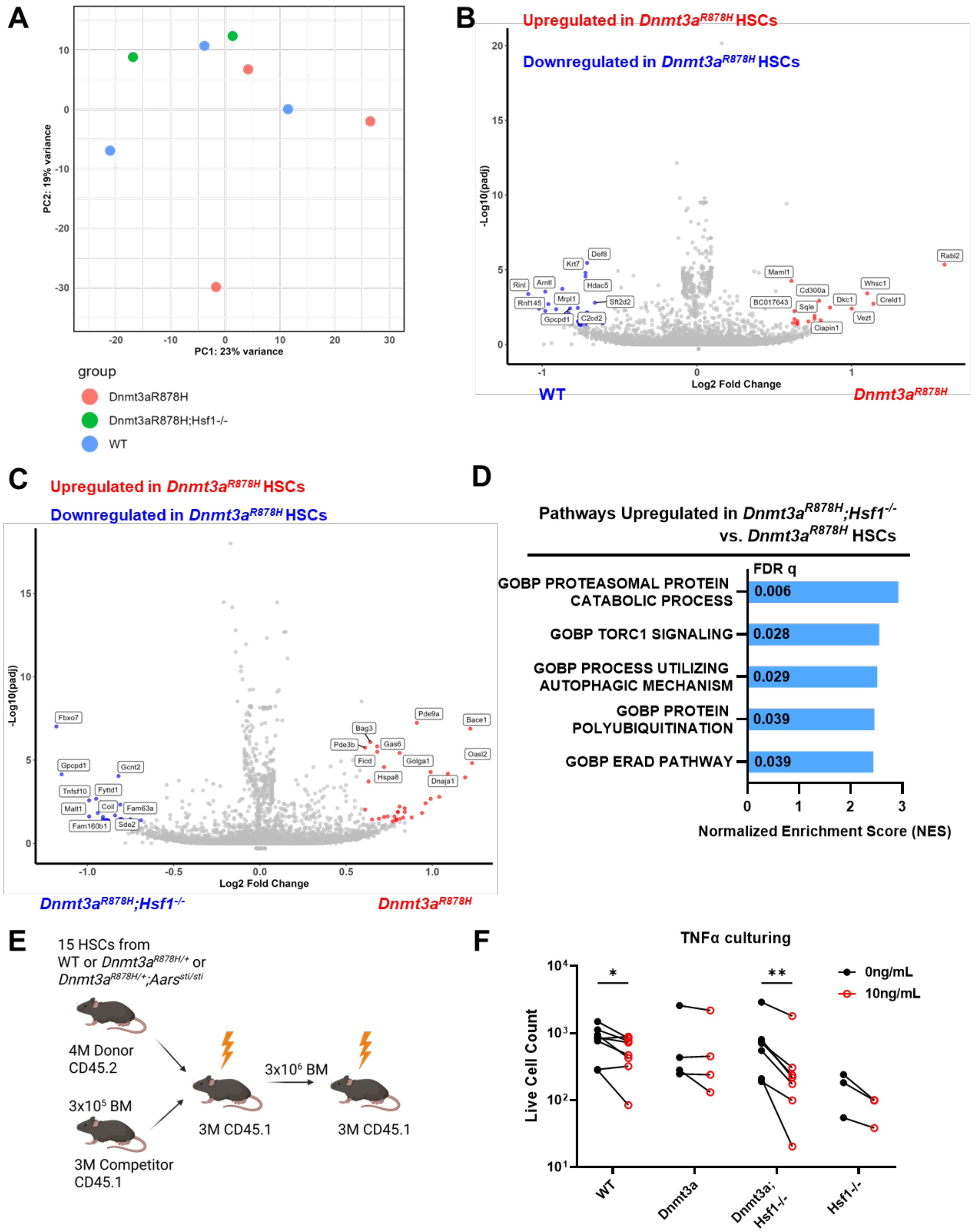
A
PC1: 23% variance
PC2: 19% variance
group
Dnmt3aR878H
Dnmt3aR878H;Hsf1-/-
WT
B
Upregulated in Dnmt3aR878H HSCs
Downregulated in Dnmt3aR878H HSCs
-Log10(padj)
Log2 Fold Change
WT
Dnmt3aR878H
C
Upregulated in Dnmt3aR878H HSCs
Downregulated in Dnmt3aR878H HSCs
-Log10(padj)
Log2 Fold Change
Dnmt3aR878H;Hsf1-/-
Dnmt3aR878H
D
Pathways Upregulated in Dnmt3aR878H;Hsf1-/- vs. Dnmt3aR878H HSCs
FDR q
GOBP PROTEASOMAL PROTEIN CATABOLIC PROCESS
0.006
GOBP TORC1 SIGNALING
0.028
GOBP PROCESS UTILIZING AUTOPHAGIC MECHANISM
0.029
GOBP PROTEIN POLYUBIQUITINATION
0.039
GOBP ERAD PATHWAY
0.039
Normalized Enrichment Score (NES)
E
15 HSCs from
WT or Dnmt3aR878H/+ or
Dnmt3aR878H/+;Aarssti/sti
4M Donor
CD45.2
3x10⁵ BM
3M Competitor
CD45.1
3x10⁶ BM
3M CD45.1
3M CD45.1
F
TNFα culturing
Live Cell Count
0ng/mL
10ng/mL
WT
Dnmt3a
Dnmt3a;
Hsf1-/-
Hsf1-/-

**Figure S6 (Related to Figure 6). Hsf1 influences gene expression and stress resistance of *Dnmt3a*$^{\mathbf{R878H}}$ HSCs.**
**(A)** PCA plot of WT, *Dnmt3a*$^{R878H}$ and *Dnmt3a*$^{R878H}$*;Hsf1*$^{-/-}$ HSCs (n=2-3 mice/genotype).
**(B,C)** Volcano plots showing upregulated and downregulated genes in *Dnmt3a*$^{R878H}$ as compared to (B) WT or (C) *Dnmt3a*$^{R878H}$*;Hsf1*$^{-/-}$ HSCs (n=2-3 mice/genotype). Genes highlighted in red and blue correspond to absolute fold change of 1.5 and $p_{adj} < 0.05$.
**(D)** Selected upregulated protein degradation-related pathways based on GSEA from RNA-sequencing in *Dnmt3a*$^{R878H}$*;Hsf1*$^{-/-}$ HSCs as compared to *Dnmt3a*$^{R878H}$ HSCs (n=2-3 mice/genotype).
**(E)** Schematic for competitive, serial transplantation of 15 4-month-old WT, *Dnmt3a*$^{R878H}$*, Dnmt3a*$^{R878H}$*;Aars*$^{sti/sti}$ HSCs into irradiated recipients. Data are shown in (Fig. 6F,G).
**(F)** Line graph of each cultured HSC sample with paired 0mg/mL and 10mg/mL TNFα conditions from (Fig. 6K). Statistical significance was assessed using a paired Student's *t*-test. *p<0.05, **p<0.01
